# Consistency of resting-state correlations between fMRI networks and EEG band-power

**DOI:** 10.1101/2024.05.22.595342

**Authors:** Marta Xavier, Inês Esteves, João Jorge, Rodolfo Abreu, Anne-Lise Giraud, Sepideh Sadaghiani, Jonathan Wirsich, Patrícia Figueiredo

## Abstract

Several simultaneous EEG-fMRI studies have aimed to identify the relationship between EEG band power and fMRI resting state networks (RSNs) to elucidate their neurobiological significance. Although common patterns have emerged, inconsistent results have also been reported. This study examines the consistency of these correlations across subjects and to understand how factors such as the hemodynamic response delay and the use of different EEG data spaces (source/scalp) influence them. Using three distinct EEG-fMRI datasets, acquired independently on 1.5T, 3T and 7T MRI scanners (comprising 42 subjects in total), we evaluate the generalizability of our findings across different acquisition conditions. We found consistent correlations between fMRI RSN and EEG band-power time-series across subjects in the three datasets studied, with systematic variations with RSN, EEG frequency-band, and HRF delay, but not with EEG space. Qualitatively, the majority of these correlations were similar across the three datasets, despite important differences in field strength, number of subjects and resting-state conditions. Our findings support consistent correlations across specific fMRI RSNs and EEG bands and highlight the importance of methodological considerations in interpreting them that may explain conflicting reports in existing literature.

## 1. Introduction

Functional brain networks are defined as a set of distributed brain regions that are functionally interconnected, contributing to specific cognitive functions with different levels of complexity. The study of these networks has been crucial for understanding healthy brain function, as well as identifying markers of abnormal brain activity in pathology. Resting-state studies have been particularly useful in characterizing functional networks because they reveal intrinsic patterns of neuronal integration without the need for a specific stimulation or task paradigm. Since these resting-state patterns have first been reported (Biswal et al., 1995; Raichle et al., 2001), several highly reproducible resting-state networks (RSNs) have been identified (Smith et al., 2009; Yeo et al., 2011) that are spatially consistent across different subjects as well as acquisition conditions. Over the years, functional magnetic resonance imaging (fMRI) has been the preferred imaging modality for characterizing RSNs because of its high spatial resolution, whole-brain coverage and non-invasiveness. Traditionally, RSNs have been identified using either independent component analysis (ICA) or seed-based linear regression or correlation analysis, both of which assess functional integration based on temporal correlations between fMRI signals across different brain regions.

Despite being widely used to study functional brain networks, fMRI suffers from low temporal specificity, which limits its ability to capture the temporal co-fluctuations that characterize neuronal coupling. Indeed, fMRI typically measures brain activity indirectly via the blood-oxygen-level-dependent (BOLD) signal, which reflects changes in blood oxygenation levels following neuronal activation by a few seconds. These changes result from neurovascular coupling mechanisms leading to localized changes in cerebral blood flow and volume associated with changes in the cerebral metabolic rate of oxygen, collectively known as the hemodynamic response. The delay of this response, together with the influence of non-neuronal sources such as heart rate and respiration, significantly confounds the identification of the true neuronal contributions of the BOLD signal (Logothetis et al., 2001). In contrast, electrophysiological methods like electroencephalography (EEG) or magnetoencephalography (MEG) directly measure neuronal activity with sub-millisecond temporal resolution, making them suitable for capturing the transient neuronal dynamics resulting from functional integration. However, these methods are limited by their low spatial specificity, partially due to the number of electrodes/sensors and the influence of volume conduction effects. The latter can cause neuronal activity from a single source to be detected in multiple scalp regions (Palva and Palva, 2012). Methods to reconstruct source activity from sparse scalp measurements have been developed and widely used (Michel and Murray, 2012), but may be impacted by differences in methodology as they attempt to solve an ill-posed inverse problem. Simultaneous EEG-fMRI studies have sought to identify electrophysiological correlates of fMRI RSNs, by leveraging the complementary characteristics of the two modalities. However, integrating EEG and fMRI data remains a challenge, and multiple multimodal data integration methods have been employed in the literature (Jorge et al., 2014; Abreu et al., 2018; Chang and Chen, 2021; Wirsich et al., 2023; Fleury et al., 2023).

Early studies predominantly used multiple regression frameworks to combine average EEG activity within a given frequency-band and from a set of mostly occipital electrodes with voxel-wise whole-brain fMRI activity (Goldman, 2002; Moosmann et al., 2003; Laufs et al., 2003a, 2003b; Gonçalves et al., 2006, Laufs et al., 2006). Most studies filtered EEG activity in the alpha frequency-band (8-12 Hz) and reported solely negative correlations. Mantini and colleagues (Mantini et al., 2007) proposed a different approach, seeking to correlate EEG activity from a range of frequency-bands with the fMRI activity of canonical RSNs obtained with ICA. They found that each RSN could be characterized by a specific combination of EEG frequency-bands. For the specific case of the default mode network (DMN), they found significant positive correlations with EEG band-specific power, including the alpha frequency-band (8-13 Hz), which was contradictory to much of the previous literature. Moreover, subsequent studies that attempted to replicate this approach found large variations between subjects leading to non-significant group results (Meyer at al., 2013). Some studies have extended this approach by preserving the spatial dimension of the EEG data. Jann and colleagues (Jann et al., 2010) estimated the covariance between band-specific power from each EEG channel and each fMRI RSN, and found clusters of frequency-bands with similar covariance topographies. Another study (Scheeringa et al., 2008) performed spatial ICA on the EEG scalp data and found negative correlations between a frontal midline theta independent component and the voxels within regions belonging to the DMN. Bowman and colleagues (Bowman et al., 2017) further highlighted the presence of spatial inhomogeneities on the fMRI side. They obtained both negative and positive correlations between the EEG alpha power and the fMRI activity of different independent components (ICs) belonging to the DMN. These results suggested that different subnetworks may have contrasting relationships with the same EEG rhythm, which could explain much of the inconsistency in previous studies examining the link between the DMN and alpha power.

A variety of other approaches have also been employed over the years to investigate the relationship between EEG-derived signals and fMRI RSNs. In terms of the type of EEG features considered, these have included EEG microstates (Britz et al., 2010; Musso et al., 2010; Rajkumar et al., 2018), EEG ICs (Yuan et al., 2012; Labounek et al., 2019) and non-linear measures of the EEG spectrum (Portnova et al., 2018). On the fMRI side, dynamic functional connectivity approaches (Tagliazucchi et al., 2012; Chang et al., 2013; Allen et al., 2018) have been employed to quantify fluctuations in the activity of RSNs over time. The integration strategy itself has also been subjected to variations, with some recent studies using machine learning-based prediction frameworks (Meir-Hasson et al., 2014; Simões et al., 2020; Abreu et al., 2021).

The methodological heterogeneities, often present in more than one aspect, have contributed to the inconsistencies in the findings reported in the literature and the difficulty of interpreting them. Most strikingly, even studies that employed similar data integration strategies have occasionally reported contradictory results - as exemplified in Table 1. This raises the question of whether inconsistencies are due mainly to methodological differences in data analysis or also to differences in data quality and acquisition setup and even between subject cohorts. A way to address this question is to employ the same data analysis methodology to different datasets, acquired from different subjects and under distinct conditions. We followed this approach and reviewed the studies that employed a comparable EEG-fMRI integration method, consisting in identifying fMRI RSNs using a data-driven method and correlating their activity with EEG spectral features such as the band-specific power. These are summarized in Table 1. Despite consistencies across studies, several inconsistencies can also be found: this includes variations in the DMN in the Sensorimotor and the Frontoparietal Networks (SMN and FPN, respectively), as previously detailed. For the SMN, although negative correlations with alpha and beta band powers are commonly observed (Mantini et al., 2007; Jann et al., 2010; Meyer et al., 2013), both negative (Mantini et al., 2007; Jann et al., 2010) and positive (Jann et al., 2010; Meyer et al., 2013) correlations extending to delta and theta bands are reported. Regarding the FPN, while one study (Meyer et al., 2013) identified negative correlations in alpha and beta bands, another (Jann et al., 2010) observed positive beta correlations and variable alpha correlations across different scalp regions.

**Table 1.**
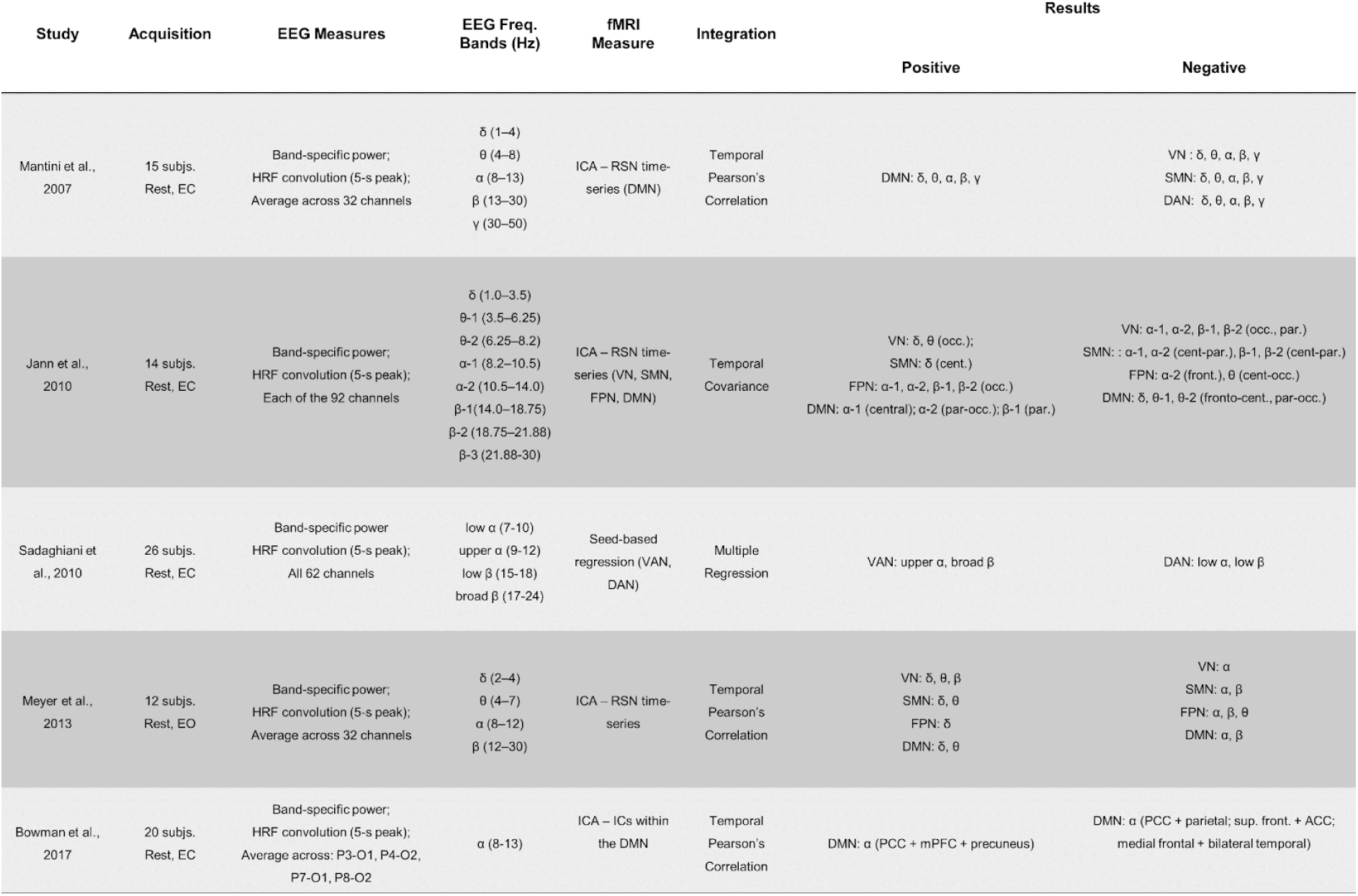
Studies that investigate the temporal correlation between fMRI-derived resting state networks (RSNs) and EEG spectral measures.

| Study | Acquisition | EEG Measures | EEG Freq. Bands (Hz) | fMRI Measure | Integration | Results |  |
| --- | --- | --- | --- | --- | --- | --- | --- |
|  |  |  |  |  |  | Positive | Negative |
| Mantini et al., 2007 | 15 subjs.<br>Rest, EC | Band-specific power;<br>HRF convolution (5-s peak);<br>Average across 32 channels | $\delta$ (1–4)<br>$\theta$ (4–8)<br>$\alpha$ (8–13)<br>$\beta$ (13–30)<br>$\gamma$ (30–50) | ICA – RSN time-series (DMN) | Temporal Pearson's Correlation | DMN: $\delta$, $\theta$, $\alpha$, $\beta$, $\gamma$ | VN: $\delta$, $\theta$, $\alpha$, $\beta$, $\gamma$<br>SMN: $\delta$, $\theta$, $\alpha$, $\beta$, $\gamma$<br>DAN: $\delta$, $\theta$, $\alpha$, $\beta$, $\gamma$ |
| Jann et al., 2010 | 14 subjs.<br>Rest, EC | Band-specific power;<br>HRF convolution (5-s peak);<br>Each of the 92 channels | $\delta$ (1.0–3.5)<br>θ-1 (3.5–6.25)<br>θ-2 (6.25–8.2)<br>α-1 (8.2–10.5)<br>α-2 (10.5–14.0)<br>β-1 (14.0–18.75)<br>β-2 (18.75–21.88)<br>β-3 (21.88–30) | ICA – RSN time-series (VN, SMN, FPN, DMN) | Temporal Covariance | VN: $\delta$, $\theta$ (occ.);<br>SMN: $\delta$ (cent.)<br>FPN: α-1, α-2, β-1, β-2 (occ.)<br>DMN: α-1 (central); α-2 (par-occ.); β-1 (par.) | VN: α-1, α-2, β-1, β-2 (occ., par.)<br>SMN: α-1, α-2 (cent-par.), β-1, β-2 (cent-par.)<br>FPN: α-2 (front.), $\theta$ (cent-occ.)<br>DMN: $\delta$, θ-1, θ-2 (fronto-cent., par-occ.) |
| Sadaghiani et al., 2010 | 26 subjs.<br>Rest, EC | Band-specific power<br>HRF convolution (5-s peak);<br>All 62 channels | low $\alpha$ (7–10)<br>upper $\alpha$ (9–12)<br>low $\beta$ (15–18)<br>broad $\beta$ (17–24) | Seed-based regression (VAN, DAN) | Multiple Regression | VAN: upper $\alpha$, broad $\beta$ | DAN: low $\alpha$, low $\beta$ |
| Meyer et al., 2013 | 12 subjs.<br>Rest, EO | Band-specific power;<br>HRF convolution (5-s peak);<br>Average across 32 channels | $\delta$ (2–4)<br>$\theta$ (4–7)<br>$\alpha$ (8–12)<br>$\beta$ (12–30) | ICA – RSN time-series | Temporal Pearson's Correlation | VN: $\delta$, $\theta$, $\beta$<br>SMN: $\delta$, $\theta$<br>FPN: $\delta$<br>DMN: $\delta$, $\theta$ | VN: $\alpha$<br>SMN: $\alpha$, $\beta$<br>FPN: $\alpha$, $\beta$, $\theta$<br>DMN: $\alpha$, $\beta$ |
| Bowman et al., 2017 | 20 subjs.<br>Rest, EC | Band-specific power;<br>HRF convolution (5-s peak);<br>Average across: P3-O1, P4-O2, P7-O1, P8-O2 | $\alpha$ (8–13) | ICA – ICs within the DMN | Temporal Pearson's Correlation | DMN: $\alpha$ (PCC + mPFC + precuneus) | DMN: $\alpha$ (PCC + parietal; sup. front. + ACC; medial frontal + bilateral temporal) |

In addressing the correlation between EEG and fMRI data, previous studies have predominantly used a canonical hemodynamic response function (HRF), which may not fully capture the known variability of the hemodynamic response across brain regions, individuals and conditions (Logothetis and Wandell, 2004). Therefore, our study expands on this approach by employing a range of HRFs to better account for and investigate the impact of these variations. Additionally, while most previous research has relied on scalp-measured EEG data, which is susceptible to volume conduction effects and lacks direct spatial correspondence with fMRI data, our study also considers the implications of using source EEG data despite the challenges in source localization.

Our main goal is to evaluate the consistency of temporal correlations between EEG-derived spectral features and fMRI RSN time-courses, both within and across datasets. By conducting a systematic consistency analysis, we aim to synthesise and clarify previous findings.

## 2. Materials and Methods

### 2.1. Datasets

Three previously acquired and reported simultaneous EEG-fMRI datasets were used in this study. The respective subject groups and acquisition setups are described below.

#### 2.1.1. Dataset 1 (1.5T)

Simultaneous EEG-fMRI data was acquired from 16 healthy volunteers, during one session of 10 min 48 s eyes-open resting-state (Deligianni et al., 2014). Of the 16 subjects, only 10 were selected for inclusion in this study based on the consistency of their EEG electrode placements, ensuring spatial uniformity within the dataset. Ethical approval was given by the local Research Ethics Committee (UCL Research ethics committee) and informed consent was obtained from all subjects. Subjects were asked to avoid movement, remain awake and fixate on a white cross presented on a black background. MRI data was recorded on a 1.5T MRI scanner (Siemens Avanto). fMRI was acquired with a T2∗-weighted gradient-echo echo-planar imaging (GRE-EPI) sequence with 300 volumes and the following parameters: TR/TE = 2160/30 ms, flip angle 75°, 30 slices (slice thickness 3.0 mm and 1 mm gap), field of view (FOV) = 210 × 210 x 120 mm3, and voxel size = 3.3 × 3.3 × 3.0 mm3. A T1-weighted structural image was also acquired (176 sagittal slices, voxel size =1.0 mm isotropic). Scalp EEG was recorded with two 32-channel MR-compatible amplifiers (BrainAmp MR, sampling rate 1kHz) and 63 electrodes (BrainCap MR). The electrodes were arranged according to the modified combinatorial nomenclature, referenced to FCz, and 1 ECG electrode was used. The EEG amplifiers were time-locked with the scanner clock.

#### 2.1.2. Dataset 2 (3T)

Simultaneous EEG-fMRI was acquired from 26 healthy volunteers, during three consecutive sessions of 10 min eyes-closed resting-state (Sadaghiani et al., 2010; Morillon et al., 2010). Out of the 26 subjects, only 23 were selected for inclusion in this study, specifically those for whom complete data from all three sessions were available for both EEG and fMRI. Ethical approval was given by the local Research Ethics Committee (CPP Ile-de France III) and informed consent was obtained from all subjects. Subjects were asked to avoid movement and remain awake. MRI data was recorded on a 3T MRI scanner (Siemens Tim-Trio). fMRI was acquired with a GRE-EPI sequence with 300 volumes (900 total for all sessions) and the following parameters: TR/TE = 2000/50 ms, flip angle 78°, 40 slices, FOV = 192 × 192 x 120 mm3, and voxel size = 3.0 x 3.0 x 3.0 mm3. A T1-weighted structural image was also acquired (176 sagittal slices, FOV = 256 x 256 mm2, voxel size = 1.0 mm isotropic). Scalp EEG was recorded with two 32-channel MR-compatible amplifiers (BrainAmp MR, sampling rate 5kHz) and 62 electrodes (Easycap electrode cap). The electrodes were referenced to FCz, and two additional electrodes, 1 ECG and 1 EOG, were used. The EEG amplifiers were time-locked with the scanner clock.

#### 2.1.3. Dataset 3 (7T)

Simultaneous EEG-fMRI was acquired from 9 healthy volunteers, during one session of 8 min eyes-open resting-state (Jorge et al., 2019). Ethical approval was given by the local Research Ethics Committee (CER-VD) and informed consent was obtained from all subjects. Subjects were asked to avoid movement, remain awake and fixate on a red cross presented on a gray background. MRI data was recorded on a 7T MRI scanner (Siemens Magnetom). fMRI was acquired with a simultaneous multi-slice (SMS) GRE-EPI sequence (3 × SMS and 2 × in-plane GRAPPA accelerations) with 480 volumes and the following parameters: TR/TE = 1000/25 ms, flip angle 54°, 69 slices, voxel size = 2.2 x 2.2 x 2.2 m3. An EPI image with reversed phase encoding direction was also acquired (5 volumes) in order to perform image distortion correction. A T1-weighted structural image was acquired with a 3D gradient-echo MP2RAGE sequence (160 sagittal slices, voxel size = 1.0 mm isotropic). Scalp EEG was recorded with two 32-channel MR-compatible amplifiers (BrainAmp MR, sampling rate 5kHz) and 63 electrodes (Easycap electrode cap). The electrodes were referenced to FCz, and one additional ECG electrode was also used. Four of the 64 electrodes (T7, T8, F5 and F6) were modified to serve as motion artifact sensors (Jorge et al., 2015), leaving 59 electrodes for EEG recording. The EEG amplifiers were time-locked with the scanner clock. Respiratory traces were also recorded (sampling rate 50Hz) with a respiratory belt from the physiological monitoring unit of the MRI system.

### 2.2. MRI Data Analysis

MRI data analysis was performed using tools from the FMRIB’s Software Library (FSL 6.0.2) (Smith et al., 2004).

#### 2.2.1. MRI Preprocessing

The T1-weighted structural image was first reoriented to the standard orientation and cropped to remove head and lower neck (using FSL’s tools fslreorient2std and robustfov), corrected for bias field inhomogeneities using FSL-FAST (Zhang, Y., 2001) and then brain-extracted using FSL-BET (Smith, 2002). The structural data was then co-registered to the standard template - Montreal Neurological Institute (MNI) (Collins et al. 1994). For this, a 12 degrees of freedom (DOF) linear transformation - estimated with FSL-FLIRT (Jenkinson and Smith, 2001; Jenkinson et al., 2002) - was used to initialize a non-linear transformation - estimated with FSL-FNIRT (Andersson J. L. R., 2007). FSL-FAST was used to segment the structural data into white matter (WM), cerebrospinal fluid (CSF) and gray matter (GM) tissues.

#### 2.2.2. fMRI Preprocessing

Non-brain tissue was removed using FSL-BET, motion correction was performed with FSL-MCFLIRT (Jenkinson, M., 2002) and data was high-pass filtered using a nonlinear filter with a cut-off period of 100s. Spatial smoothing using a Gaussian kernel with full width at half-maximum (FWHM) of about 1.5 times the voxel size was then performed (5 mm for 1.5T, 4 mm for 3T, 3 mm for 7T) using FSL-SUSAN (Smith and Brady, 1997).

Functional data was co-registered to the structural image (using FSL-FLIRT with 12 DOF) and co-registered to the standard MNI template by applying the estimated linear transformation, followed by the non-linear transformation described above to co-register the structural image to the standard template. The following nuisance regressors were linearly regressed out of the data as follows:

1. 24 motion realignment parameters: the 6 motion realignment parameters (RP) estimated with FSL-MCFLIRT were first temporarily high-pass filtered with the same filter used for the functional data, to avoid the reintroduction of artifactual variance in filtered frequencies. Then the following time-series expansions were computed: their temporal derivatives (estimated as the difference between the original time-series and the backward-shifted time-series), the quadratic term of these derivatives, and the temporal derivative of the quadratic term.
2. Motion outliers: motion outliers were estimated with FSL’s *fsl_motion_outliers*, using the metric DVARS (Power et al., 2012), which is computed as the root mean square intensity difference of volume N to volume N + 1, normalized by the median brain intensity and multiplied by 1000. To identify outliers the DVARS was thresholded at the 75th percentile +1.5 times the interquartile range.
3. WM and CSF time-series: WM and CSF masks were obtained by segmentation of the T1-weighted structural image using FFSl-FAST, and then transformed to functional space using the transformation matrices described above and eroded with a 2.2 mm (WM mask)/1.8 mm (CSF mask) gaussian kernel to minimize partial volume effects. The CSF mask was additionally intersected with the mask of the large ventricles in the MNI atlas, also transformed into the functional space. The average BOLD signal time-series in each of these masks was then computed.

##### Specific strategy for dataset 7T

7T data was preprocessed according to Abreu et al., 2021. Slice timing correction and motion correction were performed with FSL’s MCFLIRT, followed by B0 distortion correction with FSL-TOPUP (Andersson et al., 2003). Nuisance regression was performed with the following additional regressors: BOLD fluctuations related to cardiac and respiratory cycles (RETROICOR, Glover et al., 2000) and with changes in heart rate, and with depth and rate of respiration (Chang et al., 2009).

#### 2.2.3. Identification of fMRI RSNs

The following procedure was applied independently to each dataset, as illustrated in Fig 1 - Bottom. Seven canonical RSNs were identified by group-level probabilistic spatial ICA (sICA) of the fMRI data, followed by template matching in (Yeo et al., 2011): visual (VN), somatomotor (SMN), dorsal attention (DAN), ventral attention (VAN) — anatomically similar to the Salience (Seeley et al., 2007) and Cingulo-Opercular networks (Dosenbach et al., 2007) — limbic (LN), frontoparietal (FPN), and default mode (DMN).

**Fig. 1.**
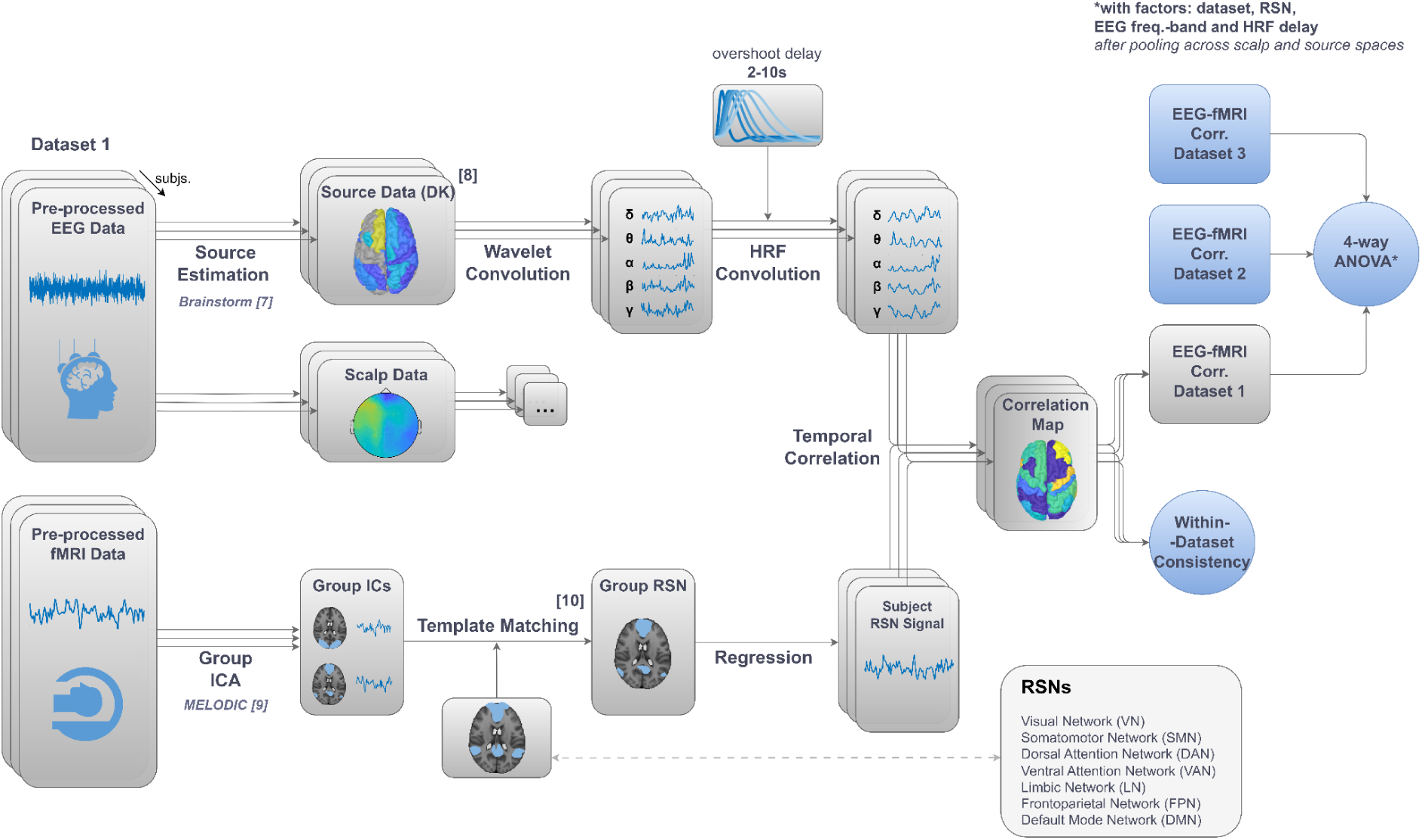
Overview of the EEG-fMRI analysis pipeline. *Bottom)* For each dataset, fMRI pre-processed data underwent group-level ICA, followed by template matching (Yeo et al., 2011), in order to identify 7 canonical RSNs. Subsequently, the resulting maps were regressed into each subject’s fMRI data to derive individual RSN time-series. ***Top)*** In parallel, EEG pre-processed data were subjected to source estimation to derive source activity data, which were spatially downsampled to align with the Desikan-Killiany (DK) atlas. Both source and scalp EEG data underwent the following analysis: estimation of the power at each frequency-band at each source node through Morlet-wavelet temporal convolution, and convolution with a family of hemodynamic response functions (HRFs) with a range of overshoot delays (2 to 10 s). ***Right)*** The resulting EEG features of each subject were temporally correlated with the simultaneously acquired fMRI RSN time-series using Pearson’s correlation. To assess between-subject consistency, t-tests against zero were conducted on spatially averaged correlation maps. Finally, a 4-way repeated measures ANOVA was performed on spatially averaged correlation values (further averaged across EEG spaces via data pooling), to evaluate the impact of dataset, RSN, EEG frequency-band, and HRF delay.

Group-level sICA was performed using FSL-MELODIC (Beckmann and Smith, 2004), whereby data from all subjects was temporally concatenated prior to estimating the ICs. The number of ICs was set in all datasets to be 30. The resulting ICs were associated with probabilistic spatial maps, consisting of independent spatial patterns common to all subjects in the dataset.

To link each of the canonical RSNs to a group independent component, IC statistical maps were thresholded (Z = 3) and binarized.. Each canonical RSN was then automatically associated with the IC yielding the highest Dice coefficient (Dice, 1945) with the respective template. The spatial maps of the selected group ICs are displayed in Fig. S1 in the supplementary material for each dataset.

To estimate subject-level time-courses for each RSN, dual regression was performed to the group-level ICs using FSL - *DualRegression* (Nickerson, 2017), and the time-courses of the respective ICs were retrieved.

### 2.3. EEG Data Analysis

#### 2.3.1. Preprocessing

Different strategies were employed for each of the datasets, as described in the original publications as indicated. These are briefly described below.

##### 1.5T and 3T

Data was preprocessed as described in Wirsich et al., 2021. EEG was corrected for the gradient artifact using template subtraction and adaptive noise cancellation. 64T-3T data was low-pass filtered at 75 Hz. Data was then downsampled at 250 Hz followed by pulse artifact correction with template subtraction, using the EEGLAB FMRIB plug-in (https://fsl.fmrib.ox.ac.uk/eeglab/fmribplugin/). For the 1.5T dataset, ICA-based denoising was performed in order to remove gradient and pulse artifact residuals as well as eye-blinks and muscle artifacts.

##### 7T

Data was preprocessed as described in Jorge et al. 2015, 2019. EEG was corrected for the gradient artifact using template subtraction. Bad-channel interpolation was performed (0-4 channels per subject, interpolated using 3-6 neighbors), followed by temporal band-pass filtering (1-70 Hz) and pulse artifact correction (using a k-means clustering approach) and downsampling at 500 Hz. Data was then corrected for motion artifacts (offline multi-channel recursive least-squares regression with the motion sensor signals), and ICA-based denoising was performed for removal of gradient and pulse artifact residuals as well as eye-blinks and muscle artifacts.

##### All datasets

Time segments contaminated with motion were identified through a semi-automatic procedure, in which time-points where the signal exceeded the mean channel time-course by 4 std were automatically pre-selected and then visually inspected (Wirsich et al., 2020). Data was bandpass-filtered at 0.5-70 Hz (1.5T), 0.3-70 Hz (3T) and 1–70 Hz (7T).

#### 2.3.2. EEG Source Imaging

EEG data was submitted to a source imaging procedure, in order to estimate the activity at the neural sources responsible for generating the recorded electrical potential distributions on the scalp. Cortical 3-dimensional surfaces (scalp/skull, skull/brain and CSF interfaces) were obtained with Freesurfer (Fischl et al., 2012, http://surfer.nmr.mgh.harvard.edu/; v7.1.0), using individual T1-weighted structural images. The remaining steps of the source imaging procedure were performed with the Brainstorm software (Tadel, 2011, http://neuroimage.usc.edu/brainstorm; version June 2022).

Individual scalp surfaces representing the head/air interface were generated based on MRI structural images. The electrode positions were co-registered into the MRI structural images by a three step procedure: initial manual adjustment of standard fiducial points, refinement by an automated algorithm, and final manual corrections to accommodate individual structural variations. Head models were estimated using individual cortical and scalp surfaces, using the OpenMEEG BEM method (Symmetric Boundary Element Method from the open source-software OpenMEEG; Gramfort et al., 2010 ; Kybic et al., 2005). 15000 constrained source dipoles were placed perpendicularly to the 3D cortical surface and a leadfield matrix was estimated to map all possible dipole configurations onto scalp potential distributions (forward problem). The minimum-norm (MN) method was applied to project scalp data onto the cortex (inverse problem), optimizing the fit using a regularizer to favor solutions with minimal brain activity amplitude.

Depth weighting was applied to modify the source covariance model, reducing dominance of shallower sources in MN current density maps. The signal covariance matrix was derived from the source model, incorporating orientation and depth weighting. Due to the absence of noise recordings, the noise covariance matrix was assumed as an identity matrix, implying uniform noise variance across sensors. These matrices were combined to form the data covariance matrix. The signal-to-noise ratio was set to 3, balancing the weight the signal model should be given relative to the noise model.

Source reconstructed data (15000 dipoles) was spatially averaged into the 68 regions of the Desikan-Killiany cortical atlas (Desikan, 2006).

#### 2.3.3. EEG Feature Extraction

For each subject, the time-courses of the following EEG features were derived: band-power (BP) in five canonical frequency-bands, using Brainstorm, as illustrated in Fig 1 - Top. Data was segmented in epochs with the duration of one fMRI TR. All features were estimated both in the scalp and source EEG spaces, for each channel (scalp) or atlas region (source) and for each epoch.

##### EEG band-power

Time-frequency (TF) decomposition was performed by temporal convolution with complex Morlet wavelets (time resolution of 3 seconds at central frequency 1 Hz). The relative power of the signal at each frequency and time-point was calculated as the square amplitude of the complex wavelet coefficients, averaged across the canonical EEG frequency-bands (delta (2-4Hz), theta (5-7Hz), alpha (8-12Hz), beta (15-29Hz), gamma (30-60Hz)) and normalized by the total power (1-60 Hz). The resulting EEG time-series were downsampled to the fMRI TR frequency, using a finite impulse response (FIR) anti-aliasing low-pass filter.

##### Hemodynamic Response Function

To account for the delay of the BOLD signal relative to the EEG, EEG features were nonlinearly transformed through convolution with a 32-seconds canonical HRF (defined as the combination of two gamma functions, one modeling the response peak and the other the post-stimulus undershoot), using the MATLAB toolbox SPM12 (Penny et al., 2007). Due to the known variability of the hemodynamic response across subjects and brain regions, each EEG feature was convolved with a family of HRFs, with varying shapes characterized by different overshoot delays (relative to onset): 2, 4, 5, 6, 8 and 10s. To ensure a coherent and physiologically plausible HRF shape across different overshoot delays, the corresponding shape parameters – undershoot delay, dispersion of both overshoot and undershoot, and their ratio – were linearly scaled in relation to the overshoot delay, preserving the dynamics of the hemodynamic response.

### 2.4. EEG-fMRI Analysis

#### 2.4.1. Joint Motion Scrubbing

Both EEG epochs and fMRI volumes that were too contaminated with motion were excluded from the analysis. The fMRI volumes discarded corresponded to the motion outliers identified with FSL’s *fsl_motion_outliers*, using the criteria described above. For the EEG, epochs were discarded according to the procedure described in Wirsich et al., 2020, whereby epochs containing motion artifacts were visually identified, after pre-selecting epochs where the signal in any channel exceeded its mean by at least 4 standard deviations.

The joint motion scrubbing procedure resulted in the following mean number of epochs discarded per subject: 10/295 (EEG = 11; fMRI = 1; overlap of EEG/fMRI = 1) for the 1.5T dataset, 114/870 (EEG = 101; fMRI = 39; overlap of EEG/fMRI = 26) for the 3T dataset, and 99/470 (EEG = 89; fMRI = 27; overlap of EEG/fMRI = 17) for the 7T dataset.

#### 2.4.2. EEG-fMRI Temporal Correlations

To estimate the degree of co-fluctuation of each fMRI RSN time-series with each EEG feature time-series, the Pearson’s correlation coefficient was computed between these signals, as illustrated in Fig 1 - Right. For each subject, fMRI RSN, EEG space, EEG frequency-band, and HRF overshoot delay, a correlation map was obtained, in which EEG-fMRI correlations are displayed at each channel (scalp space) or Desikan node (source space). Grand-mean dataset correlation maps were also obtained by averaging the subject-specific correlation maps across all the subjects of each dataset.

#### 2.4.3. Statistical analyses

To assess the consistency of the correlations obtained across subjects, two sided t-tests were conducted against a null hypothesis of zero on the spatially averaged correlation maps. Averaging correlation values across channels (in scalp space) or Desikan atlas regions (in source space) was employed as a strategy for dimensionality reduction. This decision was supported by the observation that correlations did not in general display polarized patterns (i.e., exhibiting both positive and negative values across different scalp or source areas). Prior to conducting t-tests, the normality of the data was verified using the Shapiro-Wilk test. To avoid false positives, the significance p-value threshold was adjusted for multiple comparisons (across RSNs, frequency-bands and HRF delays) by employing the False Discovery Rate (FDR) correction. Both corrected and uncorrected effects are reported.

To evaluate the impact of various factors on the spatially averaged EEG-fMRI correlation values, a 5-way repeated measures ANOVA was conducted using JASP (available at https://jasp-stats.org/). The factors included in the analysis were dataset, RSN, EEG space (source/scalp), EEG frequency-band, and HRF delay, with the spatially averaged EEG-fMRI correlation values used as the dependent variable and subjects being treated as a random factor. Significant effects identified in the ANOVA were further explored using post hoc tests, with Bonferroni correction to adjust for multiple comparisons, in order to identify specific differences among the levels of significant factors or interactions. Since no significant main effect or interactions (p>0.05) were found for the EEG space, a data pooling strategy was implemented by averaging scalp and source data. This was performed in order to preserve data from both types of EEG spaces, enhancing the generalizability of the findings. Hence, a 4-way repeated measures ANOVA was then performed with dataset, RSN, EEG frequency-band, and HRF delay as independent variables.

#### 2.4.5. Complementary analyses

Given the inherent increase in the statistical power of t-tests with the number of observations (here, the number of subjects) it is crucial to examine its effects on the presented results. Indeed, the investigated datasets exhibit significant differences in the number of subjects, with 10 subjects in the 1.5T dataset, 23 in the 3T dataset, and 9 in the 7T dataset, which could potentially impact the estimated consistency of EEG-fMRI correlations. Moreover, the datasets also present substantial variations in scan duration (10 minutes for the 1.5T dataset, 30 minutes for the 3T dataset, and 8 minutes for the 7T dataset). While this does not directly impact the t-test statistical power, it may nevertheless influence the robustness of the temporal correlation estimation given its dependence on the number of time points. Hence, complementary analyses were performed in order to test the effect of varying the sample size and scan duration on the consistency of EEG-fMRI correlations. To do so, for each dataset, segments of data were randomly selected (ranging from 1 to n consecutive minutes, over 5000 iterations) prior to computing the temporal correlations and t-stat values.

The analysis of EEG-fMRI correlations detailed in the sections above focused on a set of HRF delays from 2 to 10 seconds, equating to a −4 to +4 seconds variation around the canonical 6-s delay, typically acknowledged as physiologically relevant (Logothetis and Wandell, 2004). However, exploring a broader spectrum of HRF delays could uncover additional dynamics and lags in EEG-fMRI interactions, in particular regarding the variability of hemodynamic response across different networks and frequency-bands. As such, additional EEG-fMRI correlation analyses were conducted, extending the range of HRF delays considered from 0 to 20 seconds.

## 3. Results

### 3.1. EEG-fMRI Correlations

The spatial maps of EEG-fMRI correlations, obtained for each fMRI RSN and EEG frequency-band, in both scalp and source spaces, are presented in Fig. 2 (for the canonical HRF delay of 6s) and in Figs. S2-S6 (for the HRF delays of 2s, 4s, 5s, 8s and 10s), averaged across subjects for each dataset. In Fig 3, the corresponding spatially averaged correlations obtained for all HRF delays from 2s to 10s, showing how the overall patterns of these correlations evolve with the EEG to fMRI delay considered.

**Fig. 2.**
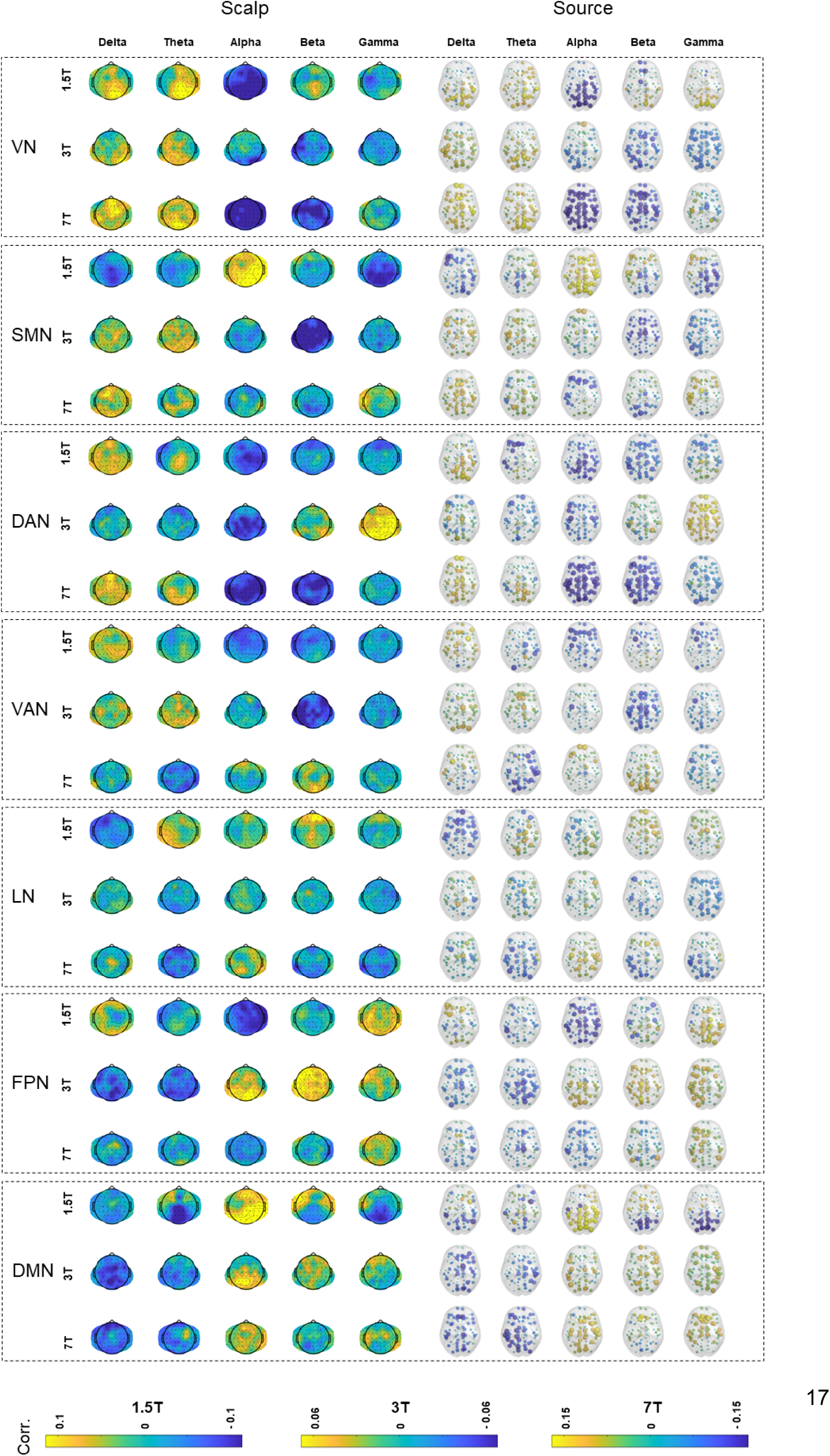
Spatial maps of EEG-fMRI correlations. The subject-averaged spatial maps of the temporal correlations, obtained with the canonical HRF with a 6-s overshoot delay, are presented for each fMRI RSN (rows) and EEG frequency-band in both scalp and source spaces (columns), for the three datasets (1.5T, 3T, and 7T).

**Fig. 3.**
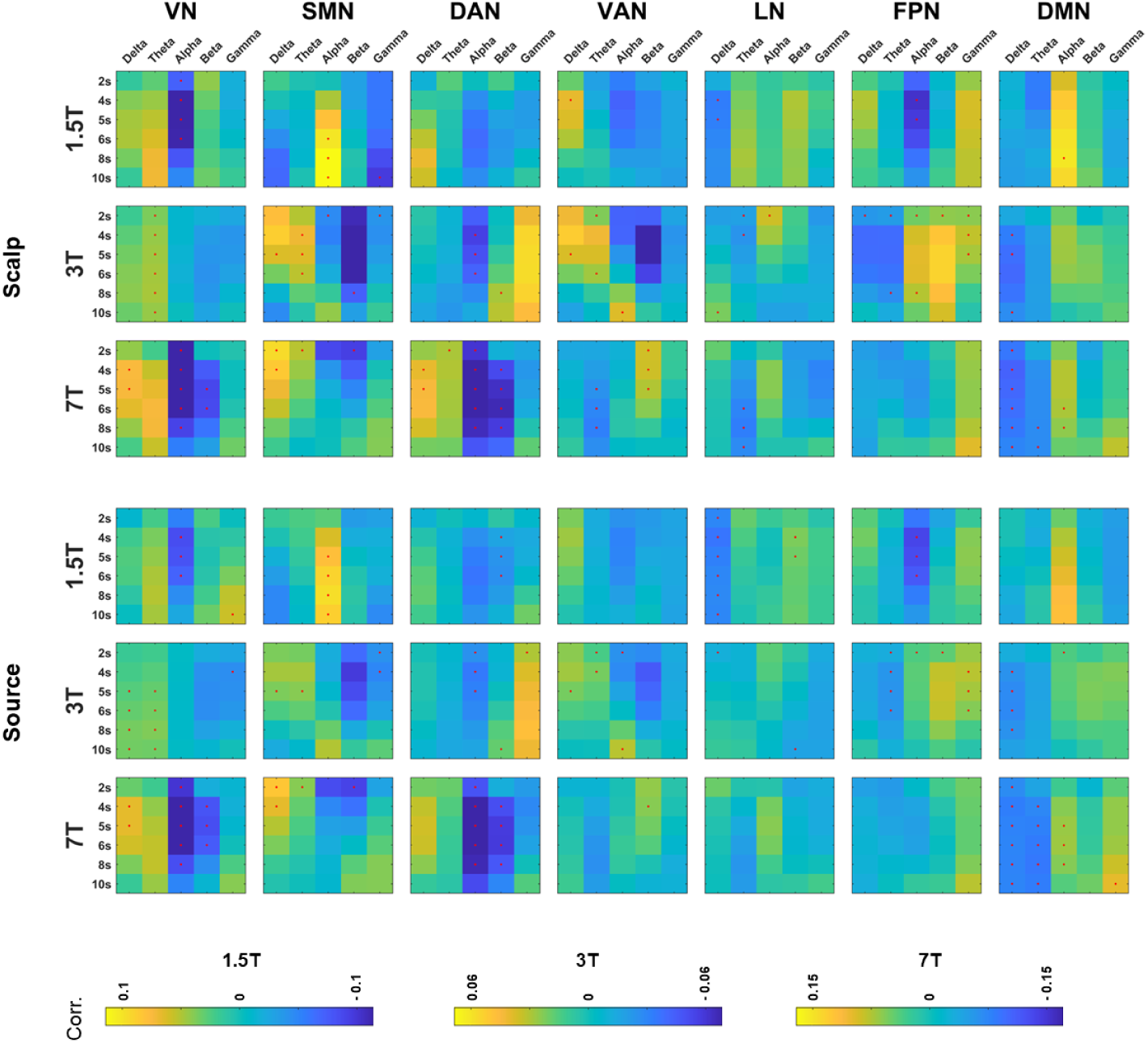
EEG-fMRI correlations across HRF delays. The heatmaps display the value of the spatially averaged EEG-fMRI correlations, averaged across subjects and for each EEG space (scalp/source), for different HRF delays: 2, 4, 5, 6,8 and 10s. Each subplot corresponds to a combination of the independently acquired datasets (1.5T, 3T and 7T) and 7 canonical RSNs. Red dots denote correlation values that are significantly different from zero, as per t-tests (uncorrected for multiple comparisons)

#### 3.1.1. Consistency across subjects

EEG-fMRI correlations, averaged across subjects, channels / regions and spaces (scalp and source) of the temporal correlations, obtained for each dataset, for each fMRI RSN, EEG frequency-band and HRF delay, are presented in Fig. 4. Fig. S7 of the supplementary material provides separate results for scalp and Desikan-Killiany data. Several significant correlations were identified between fMRI RSNs and EEG band-power for specific HRF delays, revealing complex interactions between RSN, frequency-band and HRF delay.

**Fig. 4.**
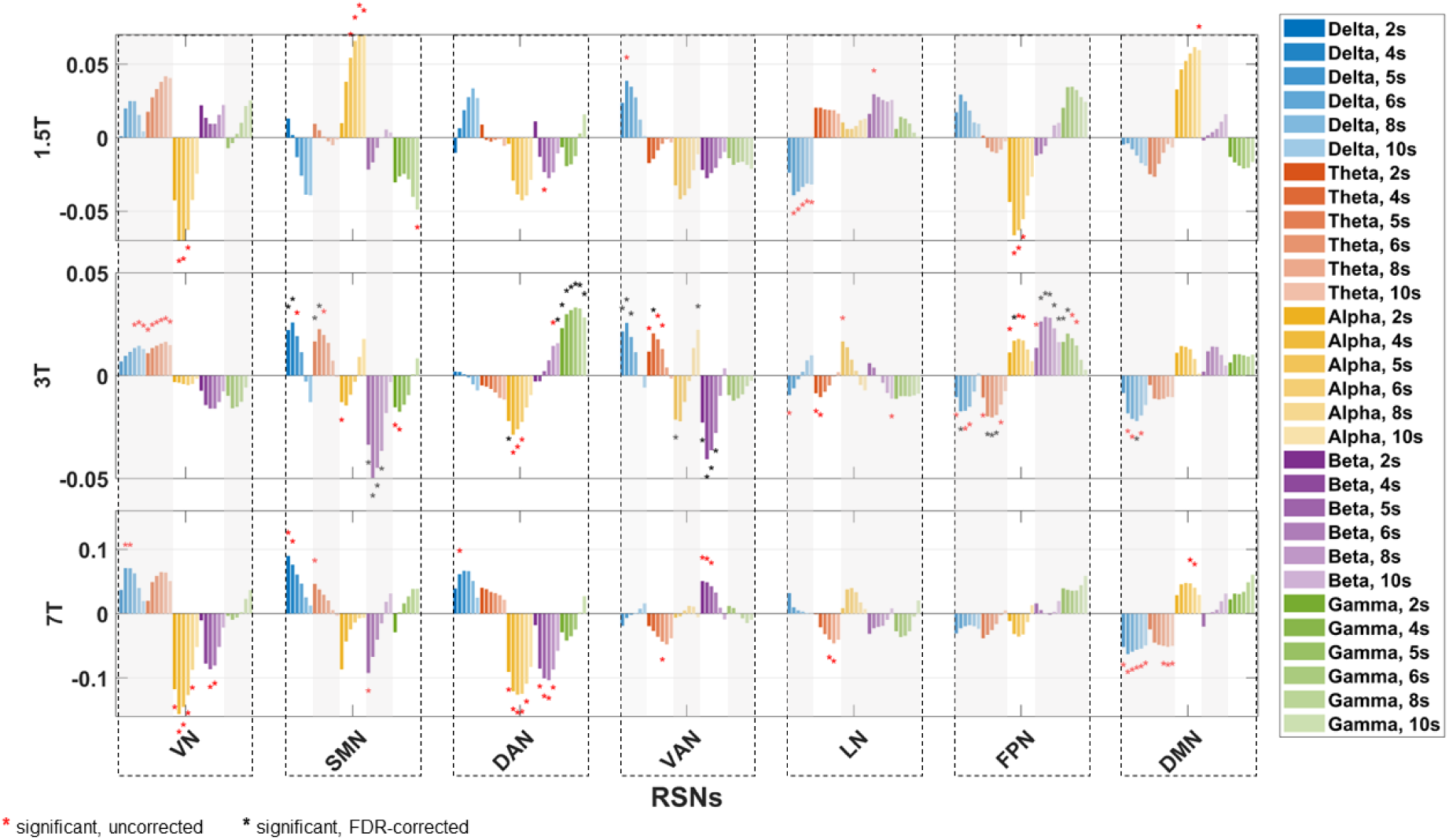
Spatially Averaged EEG-fMRI Correlations. The bar plots depict the average across subjects, channels / regions and spaces (scalp and source) of the temporal correlations, obtained for each fMRI RSN (columns), EEG frequency-band (colors) and HRF delay (hues), for each dataset (rows). Data were pooled from both scalp and source (Desikan-Killiany atlas) EEG spaces to derive average correlation values. Asterisks denote correlations significantly different from zero at p<0.05, as determined by a t-test across subjects; red and black asterisks represent uncorrected and False Discovery Rate-corrected results, respectively. Grey shading highlights RSN / frequency-band pairs that did not exhibit significant differences in correlation (pooled across delays) between datasets, according to the post-hoc analysis of the interaction between dataset, RSN, and frequency-band obtained with the 4-way repeated measures ANOVA (p<0.05, Bonferroni-corrected).

#### 3.2.2. Interactions between fMRI RSNs, EEG frequency-bands and HRF delays

The 4-way repeated measures ANOVA revealed significant main effects for dataset (F=7.7, p < 0.001), RSN (F=5.0, p<0.001), frequency-band (F=25.5, p<0.001), and HRF delay (F=3.1, p=0.008). In terms of pairwise interactions, significant findings were observed between dataset and RSN (F=5.3, p<0.001), dataset and frequency-band (F=14.8, p<0.001), RSN and frequency-band (F=39.2, p<0.001), and frequency-band and HRF delay (F=2.5, p<0.001). Additionally, a significant triple interaction was identified between dataset, RSN, and frequency-band (F=17.8, p<0.001). No significant quadruple interactions were found.

##### Interaction between dataset, RSN, and frequency-band

Post-hoc analyses, adjusted with Bonferroni correction for a family of 105, were conducted to explore the significant (p<0.05) 3-way interaction identified between dataset, RSN, and frequency-band. These effects are reported below for each network separately.

##### Visual Network

Consistent positive correlations were noted across all datasets in the delta and theta frequency-bands, with no significant differences identified between them. For the alpha band, correlations were negative across all datasets, with a notable near-zero correlation in the 3T dataset; these correlations were found to be significantly different across datasets. Variations were also observed in beta band correlations: while the 1.5T dataset demonstrated positive correlations, both the 3T and 7T datasets presented negative correlations, with these differences being statistically significant. Although gamma band correlations were not significantly different across datasets, they displayed a nuanced pattern: the 3T dataset showed slightly negative correlations, whereas the 1.5T and 7T datasets revealed slightly negative correlations at lower HRF delays and slightly positive correlations at higher HRF delays.

##### Somatomotor Network

Correlations in the delta and alpha bands demonstrated significant differences between datasets. Specifically, the 1.5T dataset exhibited negative correlations in the delta band and positive correlations in the alpha band, contrasting with the other datasets that showed positive and negative correlations in the delta and alpha bands, respectively. On the other hand, beta band correlations consistently presented as negative across all datasets, and theta band correlations as predominantly positive, both without significant differences between datasets. A significant variability was detected in the gamma band correlations among datasets: negative for the 1.5T dataset, predominantly negative for the 3T dataset, and predominantly positive for the 7T dataset.

##### Dorsal Attention Network

Correlations in all bands exhibited significant differences between datasets. The delta band showed predominantly positive correlations for the 1.5T and 7T datasets and was near zero for the 3T. Theta band correlations were close to zero for 1.5T, slightly negative for 3T, and slightly positive for 7T. Alpha band correlations were notably negative across all datasets, while beta band correlations were mostly negative, with an exception for higher HRF delays in the 3T dataset. Lastly, gamma band correlations were distinctly positive for the 3T and slightly negative for the other datasets, except for higher HRF delays.

##### Ventral Attention Network

Correlations in the delta band did not significantly differ between datasets, demonstrating positive correlations for the 1.5T and 3T datasets and being close to zero for the 7T dataset. Contrarily, theta band correlations were significantly different, presenting positive values for the 3T dataset and negative for the other two. Alpha band correlations showed a nuanced profile: negative for the 1.5T dataset, around zero for the 7T dataset, and transitioning from highly negative at lower HRF delays to highly positive at higher HRF delays in the 3T dataset. Beta band correlations were negatively associated in the 1.5T and 3T datasets and positive in the 7T dataset. Lastly, gamma band correlations, largely negative or near zero, did not exhibit significant differences across datasets.

##### Limbic Network

Correlations were mostly negative in the delta band, with no significant differences across datasets. The theta band showed a significant difference between datasets, exhibiting positive correlations in the 1.5T dataset and negative in the remaining two. Alpha band correlations were predominantly positive across all datasets without significant differences. Beta band correlations were positive in the 1.5T dataset and approximated zero in the other two, while gamma band correlations were close to zero in all three datasets, being slightly positive in the 1.5T and slightly negative in the others, with no significant differences observed between them.

##### Frontoparietal Network

Alpha band was the only frequency-band with significant differences between datasets: highly negative for the 1.5T dataset, highly positive for the 3T dataset, and slightly negative for the 7T dataset. Although not yielding significant differences, the delta band displayed negative correlations for the 3T and 7T datasets and positive for the 1.5T. Furthermore, the theta band was consistently negative across all datasets, and both beta and gamma bands presented primarily positive correlations across the datasets.

##### Default Mode Network

Despite showing significant differences in correlation values, consistent correlations across all three datasets were noted for delta, theta, alpha, and beta frequency-bands for nearly all HRF delays. Specifically, correlations in delta and theta bands were negative, while alpha and beta bands were positive (with an exception in the beta band at a 2-second delay for the 7T dataset, presenting a slightly negative yet nearly zero correlation). For the gamma band, correlations were positive in the 3T and 7T datasets and negative in the 7T dataset, being significantly different across datasets.

##### Interaction between frequency-band and HRF delay

Post-hoc analyses, adjusted with Bonferroni correction for a family of 30, were conducted to explore the significant (p<0.05) two-way interaction identified between frequency-bands and HRF delays. Specifically, for the 2s delay, correlations in both the delta and theta bands were significantly higher than in the alpha band. Similar patterns were noted for the 4s and 5s delays, where delta and theta bands consistently exhibited higher correlations than the alpha and beta bands. At the 6s delay, the delta band again yielded significantly higher correlations than the alpha band. No significant differences were identified within each band across the various delays.

### 3.2. Complementary analyses

#### 3.2.1. Effect of the Number of Subjects and Scan Duration

Figures 5 and 6 present the effects of varying the sample size and the scan duration on the significance of EEG-fMRI temporal correlations, respectively. Figs. S8-S11 in the supplementary material present these effects separately for the scalp and source spaces.

**Fig. 5.**
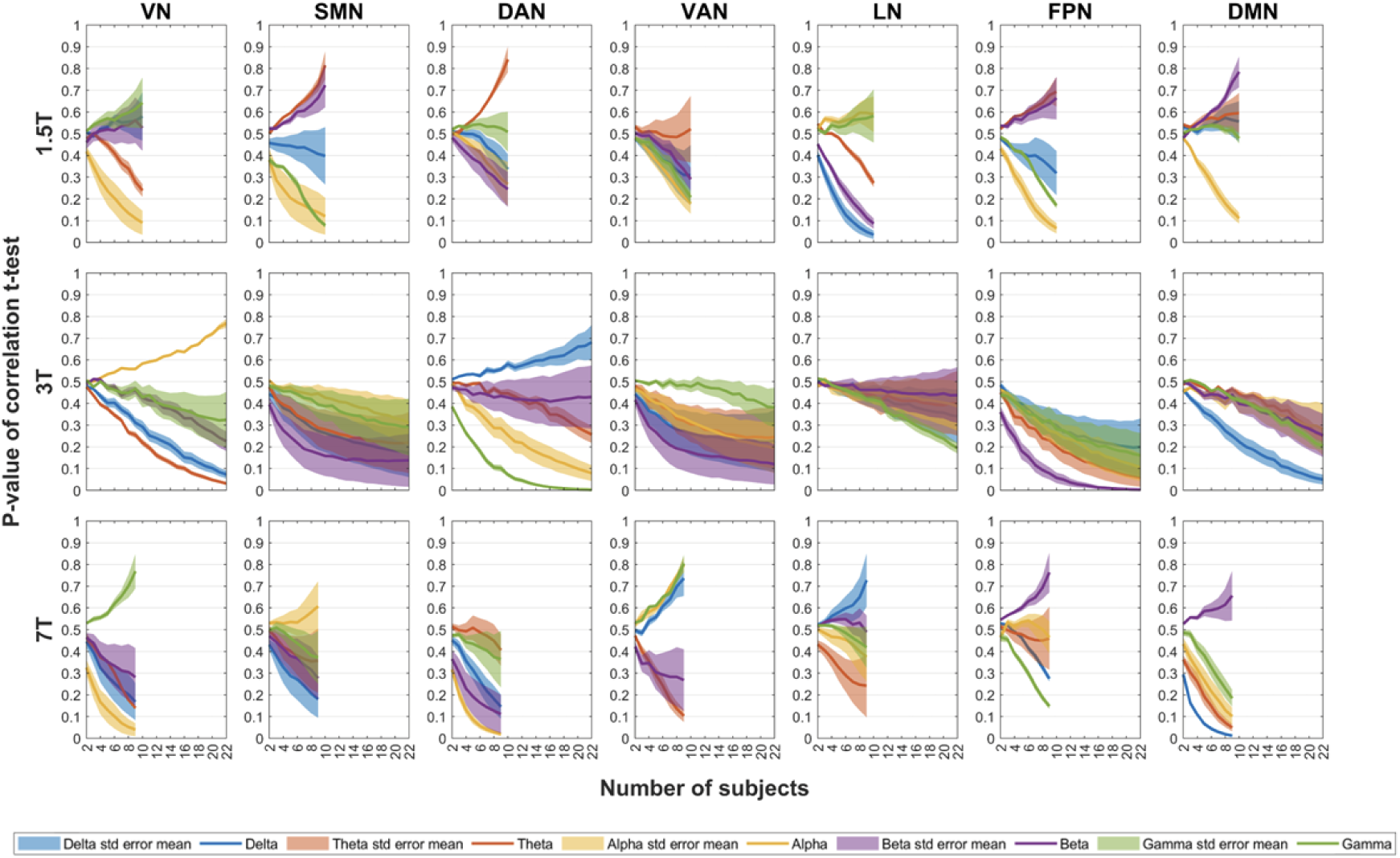
Effect of the number of subjects on the significance of EEG-fMRI correlations. Impact of increasing number of subjects on the p-value of the t-statistics derived from EEG-fMRI spatially averaged temporal correlations. Data were pooled from both scalp and source (Desikan-Killiany atlas) EEG spaces to derive average correlation values. Distinct colors denote EEG band-power across delta, theta, alpha, beta, and gamma bands, with shaded areas indicating the standard mean error across a set of HRF delays: 2s, 4s, 5s, 6s, 8s, and 10s. Rows correspond to each EEG-fMRI dataset (labeled as 1.5T, 3T, and 7T), while columns correspond to the seven canonical fMRI RSNs. For each dataset, subjects were randomly sampled (ranging from 1 to n subjects, over 5000 iterations) prior to computing the t-stat values.

**Fig. 6.**
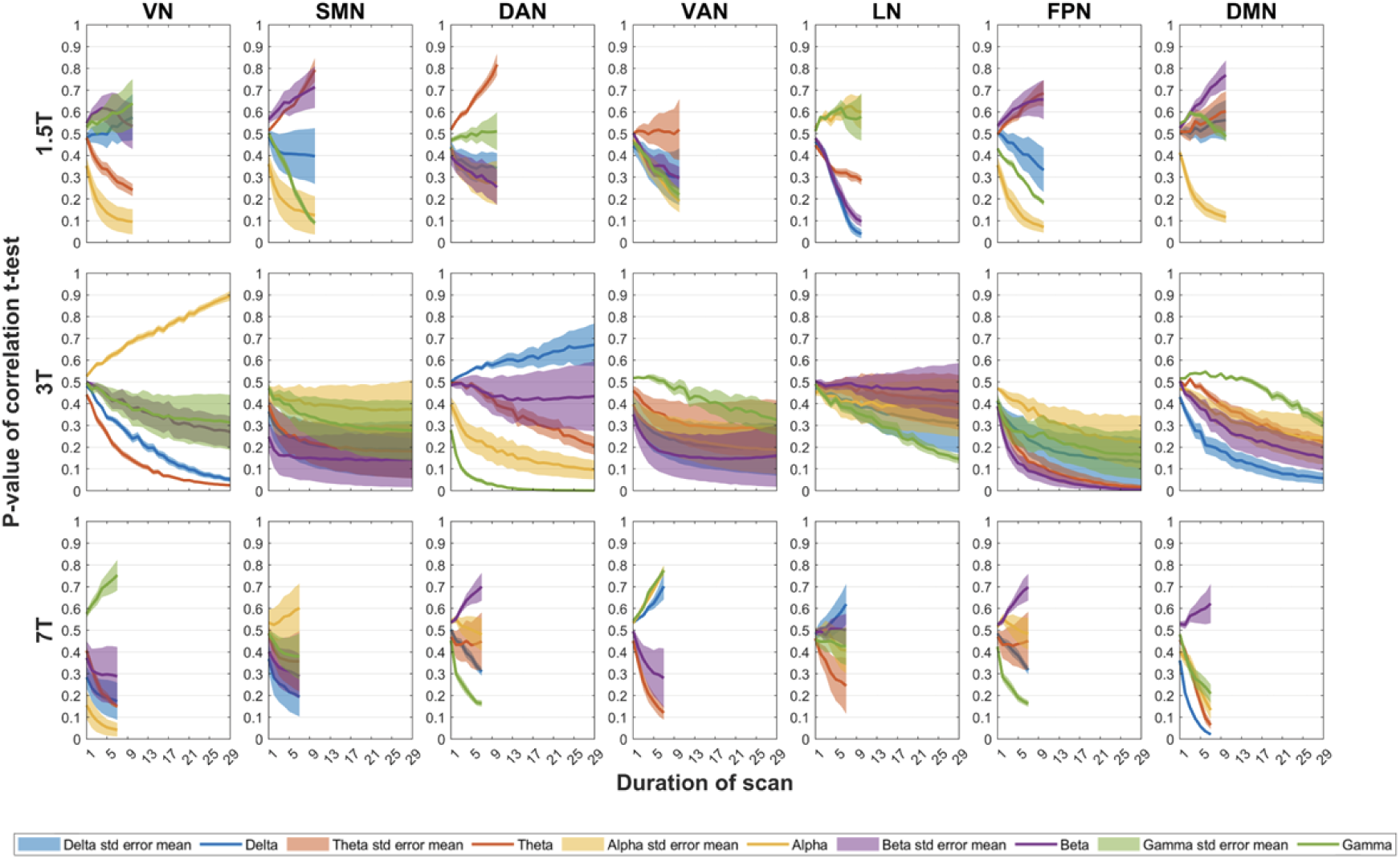
Effect of the scan duration on the significance of EEG-fMRI correlations. Impact of increasing scan duration on the p-value of the t-statistics derived from EEG-fMRI spatially averaged temporal correlations. Data were pooled from both scalp and source (Desikan-Killiany atlas) EEG spaces to derive average correlation values. Distinct colors denote EEG band-power across delta, theta, alpha, beta, and gamma bands, with shaded areas indicating the standard mean error across a set of HRF delays: 2s, 4s, 5s, 6s, 8s, and 10s. Rows correspond to each EEG-fMRI dataset (labeled as 1.5T, 3T, and 7T), while columns correspond to the seven canonical fMRI RSNs. For each dataset, segments of data were randomly selected (ranging from 1 to n consecutive minutes, over 5000 iterations) prior to computing the temporal correlations and t-stat values.

#### 3.2.2. Correlations Across Extended HRF delays

Fig. 7 explores EEG-fMRI correlations and their significance, across an extended range of HRF overshoot delays, from 0s to 20s. Fig. S12 of the supplementary material provides separate results for scalp and Desikan-Killiany data.

**Fig. 7.**
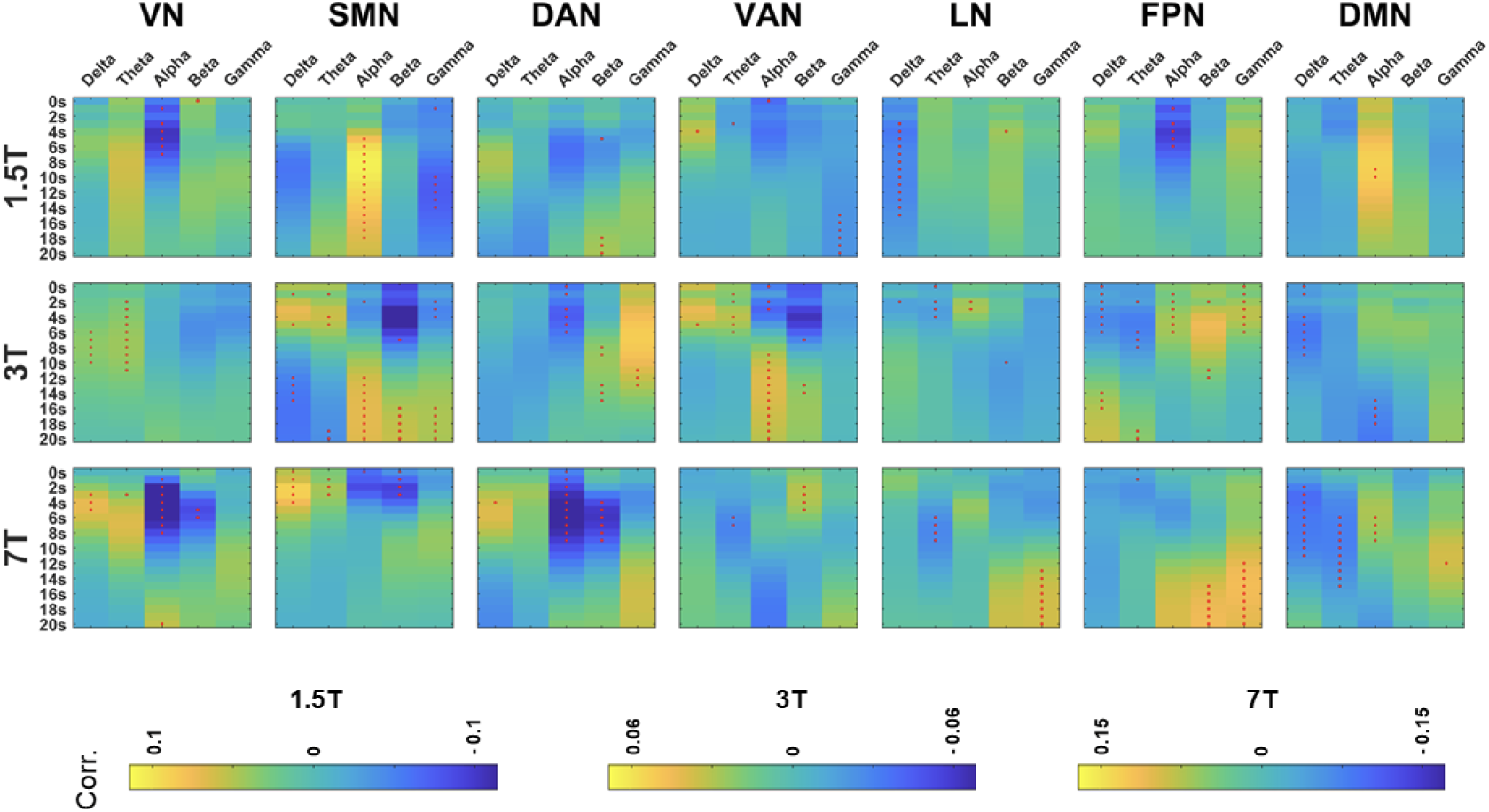
EEG-fMRI correlations across an extended range of HRF delays. The heatmaps display the value of the spatially averaged EEG-fMRI correlations, further averaged across subjects and both EEG spaces (scalp and source), for different Hemodynamic Response Function (HRF) delays, ranging from 0 to 20s. Each subplot corresponds to a combination of the independently acquired datasets (1.5T, 3T and 7T) and 7 canonical RSNs. Red dots denote correlation values that are significantly different from zero, as per t-tests (uncorrected for multiple comparisons). In each subplot, the correlation values corresponding to increasing HRF delays are displayed from top to bottom, and the values corresponding to each EEG frequency-band are displayed from left to right.

## 4. Discussion

By systematically analysing three different resting-state simultaneous EEG-fMRI datasets, we found consistent correlations across subjects between the time course of the seven canonical RSNs and the concurrent fluctuations of EEG band power in the five canonical frequency-bands. Each RSN exhibited a distinct, frequency- and delay-dependent spatial distribution of correlation with EEG power co-fluctuations. Each spatial distribution exhibited a main polarity (either positive or negative), with no evidence of opposite polarities within the same map, which prompted the use of spatially averaged correlations in subsequent statistical analyses to enhance the interpretability of results via dimensionality reduction. Significant variations were observed in the spatially averaged EEG-fMRI RSN correlations across different RSNs and EEG frequency-bands, which also significantly varied with the haemodynamic and (HRF) delays considered. Additionally, since no significant main effects or interactions were found for the EEG space (scalp vs. source) on EEG-fMRI RSN correlations, a pooling strategy was adopted in subsequent analyses, averaging both scalp and source spatially averaged correlations together.

### Relation with previous studies

Here we discuss the consistency of the EEG-fMRI correlations across the three datasets considered in our study, and in relation to previous studies, separately for each RSN.

#### Visual Network

Across datasets, the VN exhibited significant positive correlations with the delta and theta bands, as well as significant negative correlations with the beta band, aligning with previous findings (Mantini et al., 2007; Jann et al., 2010; Meyer et al., 2013). Negative correlations between the VN and the alpha band were significant in the eyes-open 1.5T and 7T datasets, while the eyes-closed 3T dataset showed non-significant negative correlations. This pattern across datasets suggests a more pronounced anti-correlation between alpha power and the VN during eyes-open resting state. Historically, alpha oscillations, especially in occipital areas, have been associated with sensory inhibition, notably increasing during eyes-closed rest and decreasing or becoming desynchronized during visual tasks or eyes-open rest, a phenomenon known as the Berger effect (Berger, 1929). This inhibitory role of alpha oscillations, often suppressed during visual attention (Fox and Snyder, 2011), might explain the observed stronger negative correlation with the VN in eyes-open conditions, acting as a mechanism to inhibit visual information processing. Additionally, these results align with numerous studies indicating a negative co-fluctuation between alpha power and fMRI signals in visual areas during resting state (Goldman et al., 2002; Moosmann et al., 2003; Laufs et al., 2006).

#### Somatomotor Network

Significant positive correlations with the delta and theta bands, alongside negative correlations with the beta and gamma bands, were identified in at least two datasets, aligning with findings from previous studies (Jann et al., 2010; Meyer et al., 2013; Mantini et al., 2007). Given that theta rhythms are often associated with sensorimotor integration (Caplan et al., 2003), these positive correlations may reflect coordinated neural activity within the SMN during resting state. In contrast, the beta central rhythm has been traditionally linked to motor control and tends to decrease during movement (Pfurtscheller, 1981); thus, the negative correlations observed suggest an inverse relationship between motor readiness and SMN BOLD activity. In fact, both beta and alpha synchronizations have been described as correlates of ’idling’ motor function, being inversely correlated with BOLD activity in the somatosensory and motor cortices (Ritter et al., 2009). It has been shown that alpha-band sensorimotor rhythm (SMR), or mu rhythm, is negatively correlated with the SMN during resting-state (Yin et al., 2016; Tsuchimoto et al., 2017). However, the correlation between the alpha band and the SMN displayed varied across datasets: while both the 3T and 7T datasets displayed the expected negative correlations (Jann et al., 2010; Meyer et al., 2013), the 1.5T dataset demonstrated significant positive correlations. Notably, DiFrancesco et al. (2008) identified positive correlations between occipital alpha EEG power and fMRI from regions within the somatomotor cortex, commenting however on the possible distinct relationship between central alpha rhythm (or μ rhythm) and occipital alpha oscillations with the BOLD activity from the somatomotor cortex. In the case of the 1.5T dataset, this cannot fully justify the discrepancy observed, since even central alpha power correlates positively with the fMRI SMN. Our deeper analysis into the dynamics of these correlations with varying HRF delays may shed light on this discrepancy. In all datasets studied, correlation patterns between alpha band and SMN evolve from near-zero/negative at shorter delays to positive/near-zero at longer delays, suggesting a biphasic hemodynamic relationship between the SMN and the alpha band, with an earlier negative inflection followed by a later positive inflection, as observed elsewhere (Prokopiou et al., 2022). In our case, the emergence of positive correlations occurs around a 6s delay in the 1.5T dataset, whereas in the other two datasets it appears only later around 10 to 12s delay. For this reason, when focusing on the canonical HRF delay of 6s, the 3T and 7T datasets exhibit the expected negative correlation but the 1.5T dataset exhibits a positive correlation. The overall patterns of correlations as a function of haemodynamic delay suggests some consistency across datasets, and with the previous literature, with a potential difference in haemodynamic delays in the 1.5T.

#### Dorsal Attention Network

The DAN showed positive correlations with the delta band in the 7T dataset and negative correlations with the alpha band in all datasets. Significant negative correlations with the beta band were found for the 1.5T and 7T dataset for HRF delays of 4-8s and significant positive correlations for longer HRF delays (8s-10s) were found for the 3T dataset. The alpha and beta band correlations are consistent with previous studies that explicitly report co-fluctuations between the fMRI DAN and EEG band-power (Mantini et al., 2007; Sadaghiani et al., 2010). Additionally, two studies (Laufs et al., 2003b; Laufs et al., 2006) also reported significant negative co-fluctuations between the alpha power and the fMRI signal in the superior parietal cortex, a region within the DAN.

#### Ventral Attention Network

The VAN showed positive correlations with the delta band for the 1.5T and 3T datasets, and with the theta band for the 3T dataset; however negative correlations with the theta band were found for the 7T dataset. In the 3T dataset, negative alpha correlations were found for the 2s HRF delay and positive alpha correlations for the 10s HRF delay. Negative beta correlations were found for the 3T dataset and positive beta correlations for the 7T dataset. Although no studies explicitly report relationships between the fMRI VAN and EEG band-power, several studies reported negative co-fluctuations between the alpha power and the fMRI signal in the inferior frontal cortex, which belongs to the VAN (Moosman et al., 2003; Laufs et al., 2003a; Laufs et al., 2006; Scheeringa, 2008). A study by Laufs et al. (2003b) reports both positive and negative co-fluctuations between the beta EEG power and the fMRI signal in the temporoparietal junction, also belonging to the VAN: positive correlations are associated with lower beta (17-23 Hz), whereas negative correlations are associated with higher beta (23-30 Hz). This dual relationship, depending on the beta frequency range considered, could potentially explain as well the divergence in our results between the 3T and 7T datasets.

#### Limbic Network

For at least two datasets, negative co-fluctuations with the delta and theta bands were found, while positive correlations were found for alpha and beta bands. Although no studies explicitly report relationships between the fMRI LN and EEG band-power, some studies reported positive co-fluctuations between the alpha EEG power and the fMRI signal in regions belonging to the limbic network: the insular cortex (Golman et al., 2002) and the anterior cingulate cortex (ACC; DiFrancesco et al., 2008).

#### Frontoparietal Network

For the 3T dataset, the FPN exhibited negative co-fluctuations with the delta and theta bands, as well as positive co-fluctuations in the beta and gamma bands, over a wide range of delays, consistently with previous studies (Mantini et al., 2007; Jann et al., 2010; Jann et al., 2010). In the alpha band, correlations were positive for the 3T dataset, but negative for the 1.5T and 7T datasets. Interestingly, an eyes-open resting-state study by Meyer et al. (2013) found negative correlations between FPN and alpha activity (similarly to what we found in the two eyes-open datasets, 1.5T and 7T), as well as positive correlations in the theta band and negative correlations in the beta band, all opposing results to the ones found for the 3T eyes-closed dataset, raising the question of weather these disparities between the three datasets could be at least partially influenced by this different condition. Notably, the eyes-closed study by Jann et al. (2010) found both positive and negative alpha correlations with the FPN, depending on the scalp region considered: positive alpha-1 (8.2-10.5 Hz) and alpha-2 (10.5-14.0 Hz) in occipital scalp electrodes and negative α-2 in frontal scalp electrodes. The topography of the scalp correlations reported in this (EC) study is in fact very similar to the one found for the 3T (EC) dataset, in which most of the positive correlations are contained in occipital channels.

#### Default Mode Network

The DMN demonstrated negative co-fluctuations with the delta and theta bands in two datasets and positive co-fluctuations with the alpha band, corroborating findings from prior research (Mantini et al., 2007; Jann et al., 2010). Notably, Scheeringa et al. (2008) identified correlations between theta power and DMN regions, such as the medial prefrontal cortex (mPFC), inferior parietal cortex, and anterior cingulate cortex (ACC). Contrasting with our findings, Mo et al. (2013) found positive correlations between the alpha power and activity within the DMN in eyes-open rest, but not in eyes-closed rest. In fact, alpha-DMN co-fluctuations are a matter of discordance in much of the existing literature, with both negative and positive correlations having been reported. Certain studies have identified negative correlations between alpha power and the DMN (Meyer et al., 2013) or DMN regions such as the ACC and inferior parietal cortex (Goldman et al., 2002; Moosmann et al., 2003; Laufs et al., 2003a; Laufs et al., 2006), while others have found positive correlations between the alpha power and DMN regions such as the ACC (DiFrancesco et al., 2008). Bowman et al. (2017) identified a dual pattern in alpha-DMN co-fluctuations, finding positive or negative correlations depending on the specific sub-network of the DMN considered. They suggest that the DMN simultaneously participates in introspective and environmental-monitoring roles, which could be reflected in both positive and negative relationships with alpha power across different regions. Similarly, Marawar et al. (2017) also found both positive and negative correlations between EEG delta and theta power and the BOLD activity within different regions of the DMN. These results highlight the complexity of the frequency modulation of the DMN, potentially explaining the varied and seemingly contradictory findings in the literature.

#### Interactions between RSNs

The DAN and DMN demonstrated inverse co-fluctuations across the delta, theta, alpha and beta bands, potentially reflecting the well-documented anti-correlation between the activity of task-positive and task-negative networks in both task and rest conditions (Fox et al., 2005; Chang et al., 2013). Traditionally, the DMN has been correlated with activity during rest and internally-oriented tasks, while the DAN has been associated with attention-demanding and externally-oriented tasks. This anti-correlation might represent a modulation in the frequency domain of brain networks, potentially competing for neural resources (Mantini et al., 2007). In alignment with this, theta power, often linked with sustained attention, exhibits positive co-fluctuations with the DAN and negative co-fluctuations with the DMN, reflecting its role in managing internal and external attention. Conversely, alpha power, which has been associated with the suppression of attention to the external environment, demonstrates negative co-fluctuations with the DAN and positive co-fluctuations with the DMN (Magosso et al., 2021).

The divergences observed in FPN co-fluctuation patterns across the 3T, 1.5T, and 7T datasets might be interpreted under a related argument. The FPN is traditionally linked to executive control and decision-making (Vincent et al., 2008), which may either relate to perceptual (externally-oriented) or introspective (internally-oriented) cognitive processes. Previous literature suggests that the FPN is functionally connected to both the DMN and the DAN (Spreng et al., 2013) and recent studies further explored this notion, investigating functional heterogeneity within the FPN (Braga et al., 2017; Dixon et al., 2018). Dixon et al. (2018) identified two main subsystems within the FPN, FPN-A and FPN-B, evident in task performance and resting state, with distinct roles in executive control. The former, functionally connected to the DMN, was theorized to be activated during internally directed attention. The latter, functionally connected to the DAN, was proposed to participate mainly in perceptual attention, facilitating interactions with the environment. In this context, our findings suggest that the FPN configuration in the 3T dataset might predominantly reflect the characteristics of the FPN-A subsystem, presenting mostly positive alpha co-fluctuations and negative theta co-fluctuations, similarly to the DMN. In contrast, the FPNs in the 1.5T and 7T datasets seem to align with the FPN-B subsystem, showing mostly negative alpha co-fluctuations, similarly to the DAN. An additional hypothesis considers the role of eyes-open versus eyes-closed condition in potentially modulating the dominant FPN mode. The eyes-open condition might promote a state of latent external attention, thereby possibly enhancing perceptual cognition. This condition could influence alpha co-fluctuations with the FPN, reflecting a subtle continuous engagement with the external stimuli, consistent with results from the eyes-open datasets. Nonetheless, confirming this hypothesis requires more detailed analysis comparing frequency modulations of the FPN in eyes-open and eyes-closed conditions within the same dataset.

On the other hand, both the limbic network and the DMN exhibited often similar EEG-fMRI correlation patterns in the delta, theta, alpha and beta bands, which might be reflective of a their related roles in internal cognition, emotional processing, and memory recall during resting-state (Greicius et al. 2003, Stephani et al., 2014). Indeed, these two networks also commonly share overlapping brain regions such as the ACC and the mPFC.

Finally, given the external attention orientation of both the DAN and VAN (Fox et al., 2006), a cooperative modulation of these two networks, leading to similar co-fluctuations with the same frequency-bands, is a logical expectation and was partially observed in our results.

Another pattern that was observed was that the EEG delta and theta power often co-fluctuated with the fMRI with the same signal, and inversely to the co-fluctuations of alpha and beta power, potentially pointing toward distinct neuronal mechanisms and cognitive states associated with these different pairs of frequency-bands.

### Impact of the haemodynamic delay

We found variability in the HRF delays at which significant EEG-fMRI correlations occur, across different RSNs, EEG frequency-bands and datasets. Notably, the most significant correlations are not always found at the canonical HRF delay of 6s, adopted in most studies exploring the relationship between EEG and fMRI RSNs. This variability might stem from the well-documented heterogeneity in neurovascular coupling across distinct brain areas and experimental conditions (Logothetis and Wandell, 2004). This observation has important implications for interpreting EEG-fMRI correlation results in the literature, highlighting the need to consider possible variability in optimal HRF delays and suggesting caution when comparing findings derived from a single, canonical HRF delay. It also highlights the importance of incorporating a range of HRF delays, specifically from 2 to 10 seconds, to fully understand the temporal dynamics of the relationship between these signals.

### Impact of EEG space

The similarity in EEG-fMRI correlations between scalp and source data, specifically the lack of significant interactions between this variable and HRF delays, suggests that the neurovascular coupling underlying the observed correlations is represented similarly in both spaces regardless of their difference in spatial specificity Crucially, this observation could inform future research in the field of EEG-fMRI correlations, by suggesting that, under specific conditions, using scalp data may provide results comparable to those derived from source-estimated data, thereby offering a methodological simplification.

### Impact of sample size

T-tests against zero were used to evaluate the significance of spatially averaged correlations across subjects, providing an assessment of between-subject consistency in correlation values for each dataset. Notably, only correlations from the 3T dataset remained significant following FDR correction. Given that this dataset comprised a considerably larger sample size (23 subjects) compared to the 1.5T and 7T datasets (10 and 9 subjects, respectively), a question arose regarding the sufficiency of the number of subjects (i.e., number of observations) to achieve statistically significant results across all datasets. By performing permutation-based tests we found that the correlation p-values in general stabilized only when considering 8-12 subjects, which could be observed only in the case of the 3T dataset. This finding suggests that discrepancies in the significance of the EEG-fMRI correlations between the 3T dataset and the other two smaller datasets may arise from this sensitivity to the sample size (the number of subjects).

### Impact of scan duration

The influence of scan duration on the correlations and their respective p-values was similarly explored, given the substantial discrepancies in scan durations among the three datasets (10 min for the 1.5T dataset, 30 min for the 3T dataset, and 8 min for the 7T dataset). The hypothesis is that correlations would stabilize and become more robust with more prolonged scan durations, potentially yielding reduced variability across subjects. This hypothesis found support by finding that correlations for the 3T dataset stabilize around a scan duration of 10-15 min, which surpasses the scan durations of the other two datasets. This stabilization is mirrored as well for the correlation p-values, which, in most instances, also stabilize around those scan durations. Such effects could be attributed to the higher correlation magnitudes observed for these durations, or alternatively, to the joint effect of higher magnitudes and greater consistency between subjects.

### Limitations and Future Work

Our study has made significant progress in reconciling and elucidating some of the inconsistencies observed in the literature regarding the relationship between EEG band power and fMRI-derived RSN activity. However, several limitations warrant further discussion, particularly in the context of defining and identifying RSNs. As highlighted in the comprehensive review by Uddin et al. (2023), there is a lack of standardized methodologies for defining and identifying RSNs, leading to variable results. This variability might be one of the primary sources of inconsistency in the literature regarding EEG-fMRI correlations of RSNs.

The methods employed to define RSNs - for example, whether networks are delineated based on fixed parcellations or are data-driven, derived from clustering or spatial ICA - can significantly impact their spatial maps. Often, the parameters guiding data-driven methods are not consistently reported, adding another layer of variability when comparing findings across different studies.There is also a lack of consensus regarding the specific regions that constitute each network, and even the naming conventions for these networks can vary, particularly for networks other than the well-recognized visual, somatomotor, and default mode networks. In the context of spatial ICA, the choice of model order significantly influences the resolution of identified networks, with networks being progressively divided in sub-networks with increasing model orders. In our research, we tried to mitigate these challenges by maintaining a consistent number of components across our datasets. Despite this, variability in the spatial maps of the identified networks remained, as depicted in Figure S1 of the supplementary material. This variability could potentially account for some of the observed differences in EEG-fMRI correlation across datasets. Although not systematically analyzed in our study, it would be pertinent for future research to investigate how the variability in fMRI network spatial maps identified across datasets influences the correlations observed.

Another challenge in the consistent definition of large-scale networks concerns the interindividual variability, which highlights the importance of capturing both common and subject-specific network characteristics. The methodology used in this study, group spatial ICA followed by dual regression, provides a means to integrate group-level results with variations specific to individuals. Even so, this variability could influence the consistency of network identification across individuals, and consequently, the magnitude of the subject-averaged EEG-fMRI correlations. Future research could explore the relationship between within-subject consistency in network spatial maps and the consistency of the corresponding correlations with EEG.

Drowsiness is another potential source of variability for EEG-fMRI correlations. Alertness levels are known to vary not only throughout resting-state acquisitions but also between subjects, influencing significantly both the temporal dynamics of the EEG spectrum and fMRI resting-state networks (Makeig and Jung, 1995; Tagliazucchi and Laufs, 2014; Joliot et al., 2024). Monitoring and adjusting for varying alertness levels could help controlling for these effects (Falahpour et al., 2018).

Finally, it is important to acknowledge that functional connectivity is not a static feature but fluctuates across multiple temporal scales, even within the duration of a single scan session (Chang and Glover, 2010). Addressing this aspect requires methodologies that can capture how these time-varying dynamics contribute to the evolving patterns of the connectome over time (Keilholz et al., 2017). Additionally, understanding the implications of these dynamic changes for the correlations between EEG power and fMRI signals is crucial. Future studies could also consider the temporal evolution of these correlations, focusing on dynamic rather than static correlations to capture their time-varying relationship.

### Conclusions

Faced with the extensive literature on EEG-fMRI correlations, in particular relating EEG band-specific power to fMRI RSNs, summarizing the varied and sometimes contradictory conclusions proves challenging due to numerous methodological variations. These span from dataset discrepancies, which inherently influence results due to variations in acquisition setups and participant numbers, to differing approaches in data preprocessing and analysis methods that combine the two modalities. Our study systematically examined EEG-fMRI correlations, taking into account key parameters such as HRF delay and the space of EEG data (scalp or source), while evaluating their consistency across subjects and generalization across different datasets. These datasets varied in fMRI field strength, number of participants, scan duration, and resting-state conditions (eyes-open vs. eyes-closed). Moreover, our systematic analysis carefully explored the spatial distribution of correlation values in both EEG scalp and source spaces. Through this approach, we not only derived conclusions about EEG-fMRI RSN correlations but also highlighted the significant impact of the explored factors, providing a clear perspective to understand some of the seemingly conflicting results in existing literature. The ability to reconcile findings from different studies by considering various previously unaccounted for parameters highlights the substantial contribution of our study.

## Data and Code Availability

The 1.5T raw data is publicly available at https://osf.io/94c5t/. The other raw data will be made available by request to ALG (64Ch-3T) and JJ (64Ch-7T). The code used for data analysis is available at https://github.com/LaSEEB/eeg_fmri_consistency.

## Author Contributions

MX and PF conceived and designed the analysis. JJ, RA, ALG, SS, and JW contributed to the acquisition and curation of data. MX, IE, JJ, RA, and JW preprocessed the data. MX performed data and statistical analysis, visualization, and developed the analysis software. PF supervised the analysis. MX and PF wrote the original draft. All authors reviewed and edited the final manuscript.

## Acknowledgments

This work was supported by LARSyS funding (DOI: 10.54499/LA/P/0083/2020, 10.54499/UIDP/50009/2020 and 10.54499/UIDB/50009/2020) and PRR project Center for Responsible AI C645008882-00000055. MX was supported by the FCT doctoral grant 2021.08229.BD. JW was supported by a research position of the Faculty of Medicine, University of Geneva.

## Supplementary Material

### fMRI Resting State Network Maps

**Fig. S1.**
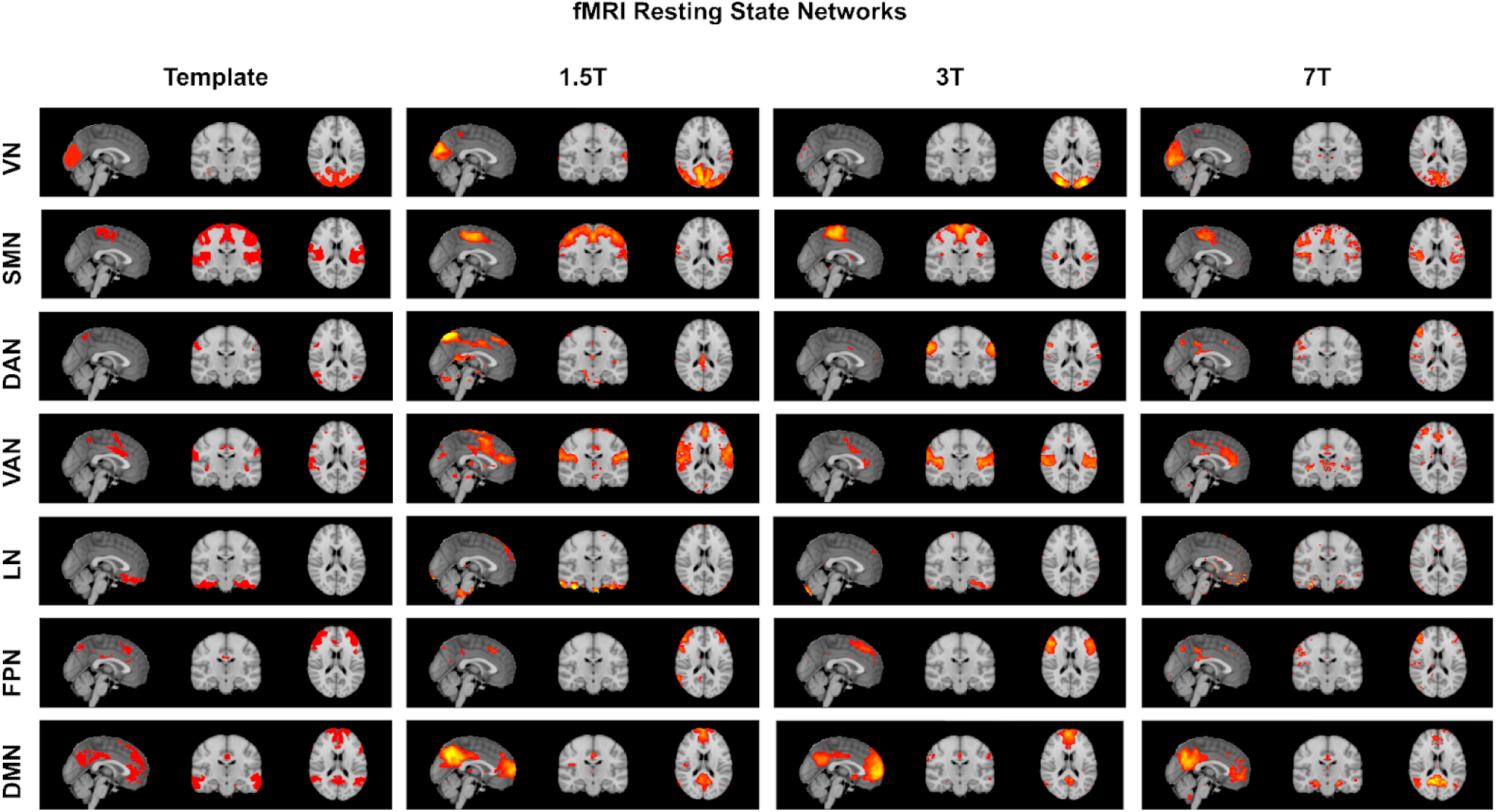
fMRI Resting State Network Maps. Spatial maps of each of the 7 canonical Resting State Networks (RSNs). From left to right: template maps (Yeo et al., 2011), group Independent Component (IC) maps obtained through group Independent Component Analysis (ICA) for three independent datasets, obtained at 1.5T, 3T and 7T, respectively. Statistical maps obtained in FSL’s FSLeyes. Sagittal, coronal and transverse views, thresholded at Z=3.

### EEG-fMRI Correlation Spatial Maps

**Fig. S2.**
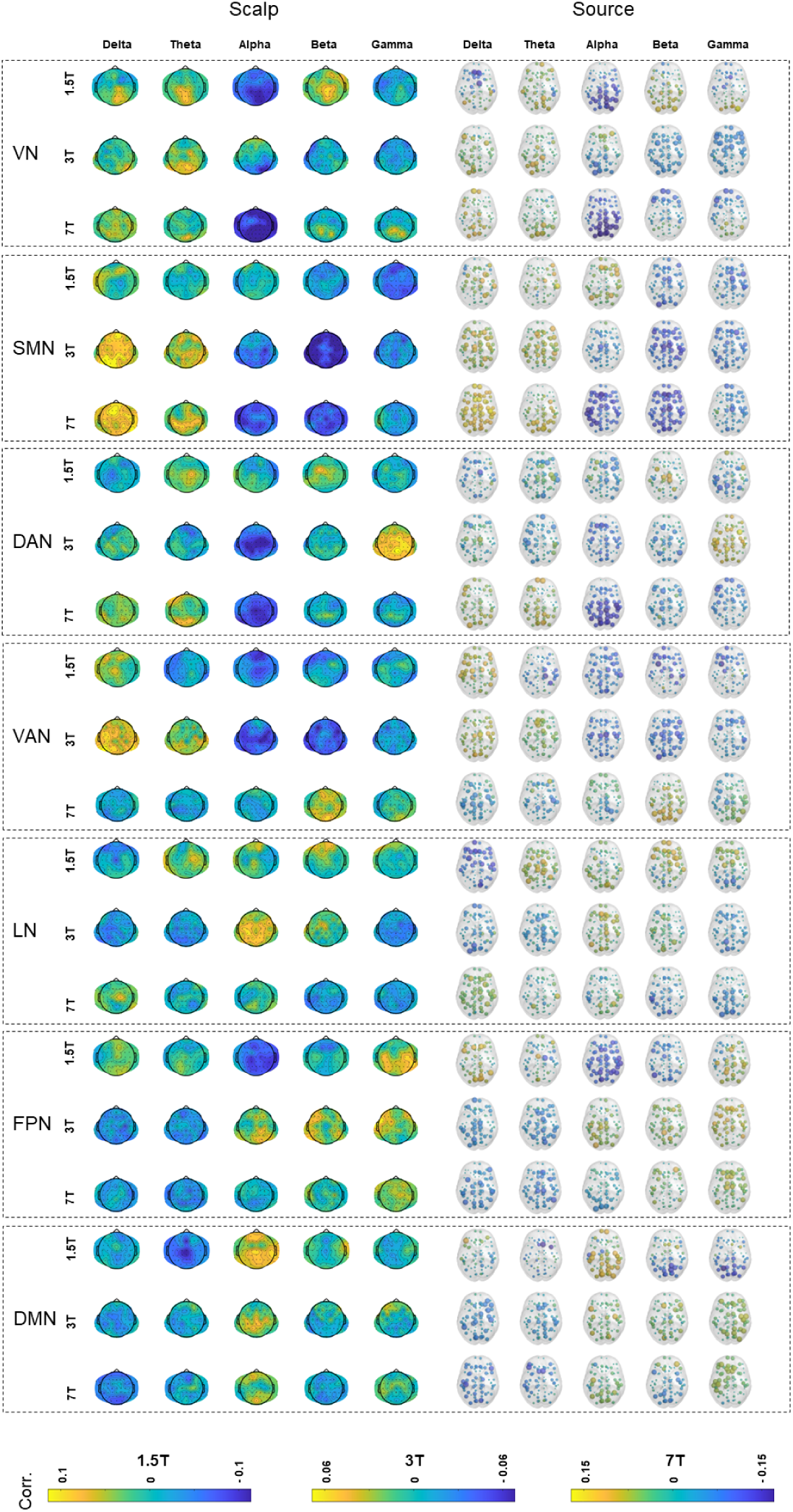
Temporal correlation spatial maps of EEG band-power and fMRI RSNs across different acquisition datasets for 2s HRF delay. The figure illustrates the subject-averaged spatial maps of the temporal correlations between 7 canonical fMRI resting state networks (RSNs) and EEG band-power across delta, theta, alpha, beta, and gamma bands for three independently acquired datasets (1.5T, 3T, and 7T), with EEG time-series convolved with an HRF with a 2-s overshoot delay. The left panel displays spatial maps related to EEG scalp data, while the right panel corresponds to source data, mapped to the Desikan-Killiany atlas.

**Fig. S3.**
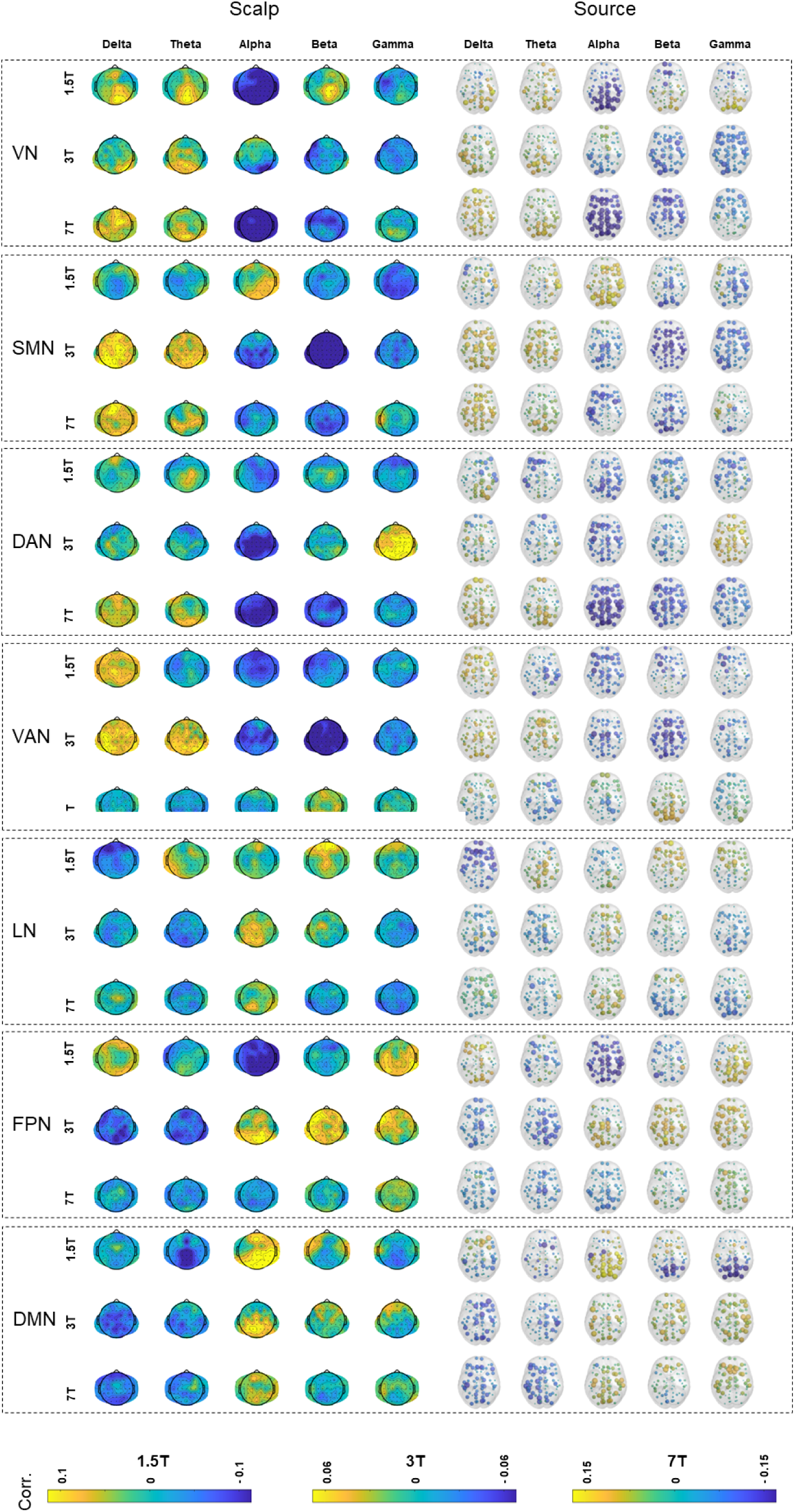
Temporal correlation spatial maps of EEG band-power and fMRI RSNs across different acquisition datasets for 4s HRF delay. The figure illustrates the subject-averaged spatial maps of the temporal correlations between 7 canonical fMRI resting state networks (RSNs) and EEG band-power across delta, theta, alpha, beta, and gamma bands for three independently acquired datasets (1.5T, 3T, and 7T), with EEG time-series convolved with an HRF with a 4-s overshoot delay. The left panel displays spatial maps related to EEG scalp data, while the right panel corresponds to source data, mapped to the Desikan-Killiany atlas.

**Fig. S4.**
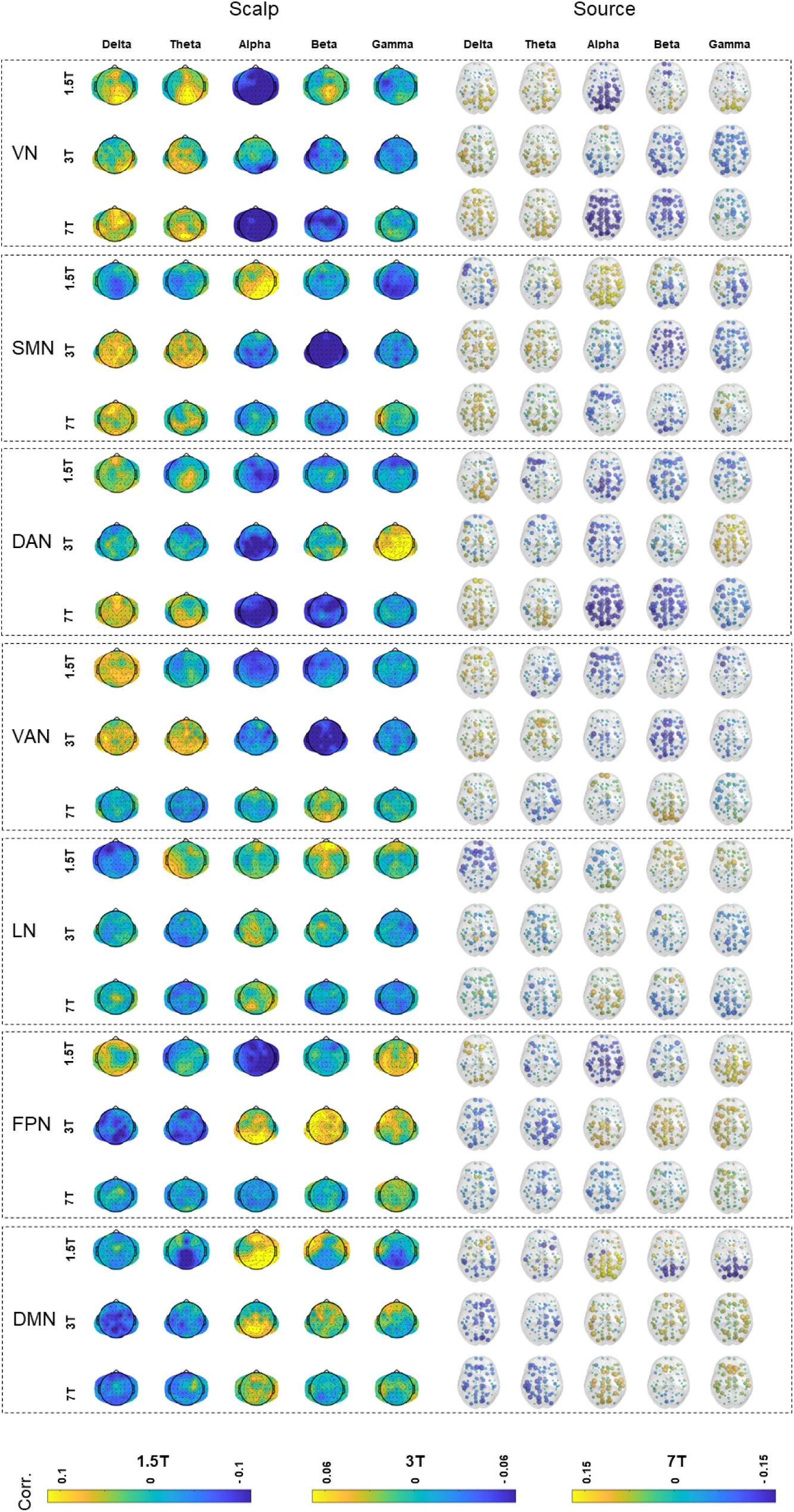
Temporal correlation spatial maps of EEG band-power and fMRI RSNs across different acquisition datasets for 5s HRF delay. The figure illustrates the subject-averaged spatial maps of the temporal correlations between 7 canonical fMRI resting state networks (RSNs) and EEG band-power across delta, theta, alpha, beta, and gamma bands for three independently acquired datasets (1.5T, 3T, and 7T), with EEG time-series convolved with an HRF with a 5-s overshoot delay. The left panel displays spatial maps related to EEG scalp data, while the right panel corresponds to source data, mapped to the Desikan-Killiany atlas.

**Fig. S5.**
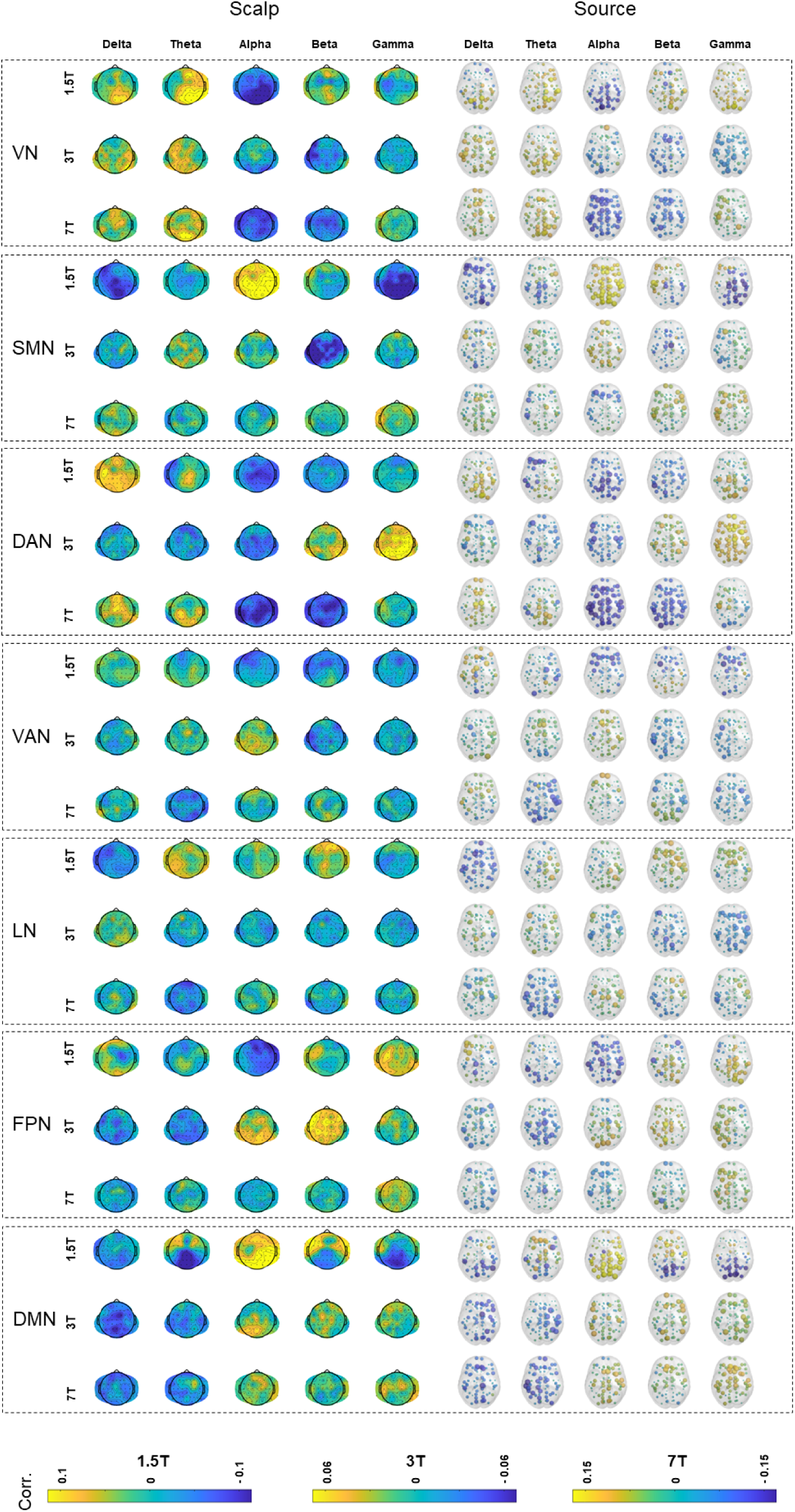
Temporal correlation spatial maps of EEG band-power and fMRI RSNs across different acquisition datasets for 8s HRF delay. The figure illustrates the subject-averaged spatial maps of the temporal correlations between 7 canonical fMRI resting state networks (RSNs) and EEG band-power across delta, theta, alpha, beta, and gamma bands for three independently acquired datasets (1.5T, 3T, and 7T), with EEG time-series convolved with an HRF with a 8-s overshoot delay. The left panel displays spatial maps related to EEG scalp data, while the right panel corresponds to source data, mapped to the Desikan-Killiany atlas.

**Fig. S6.**
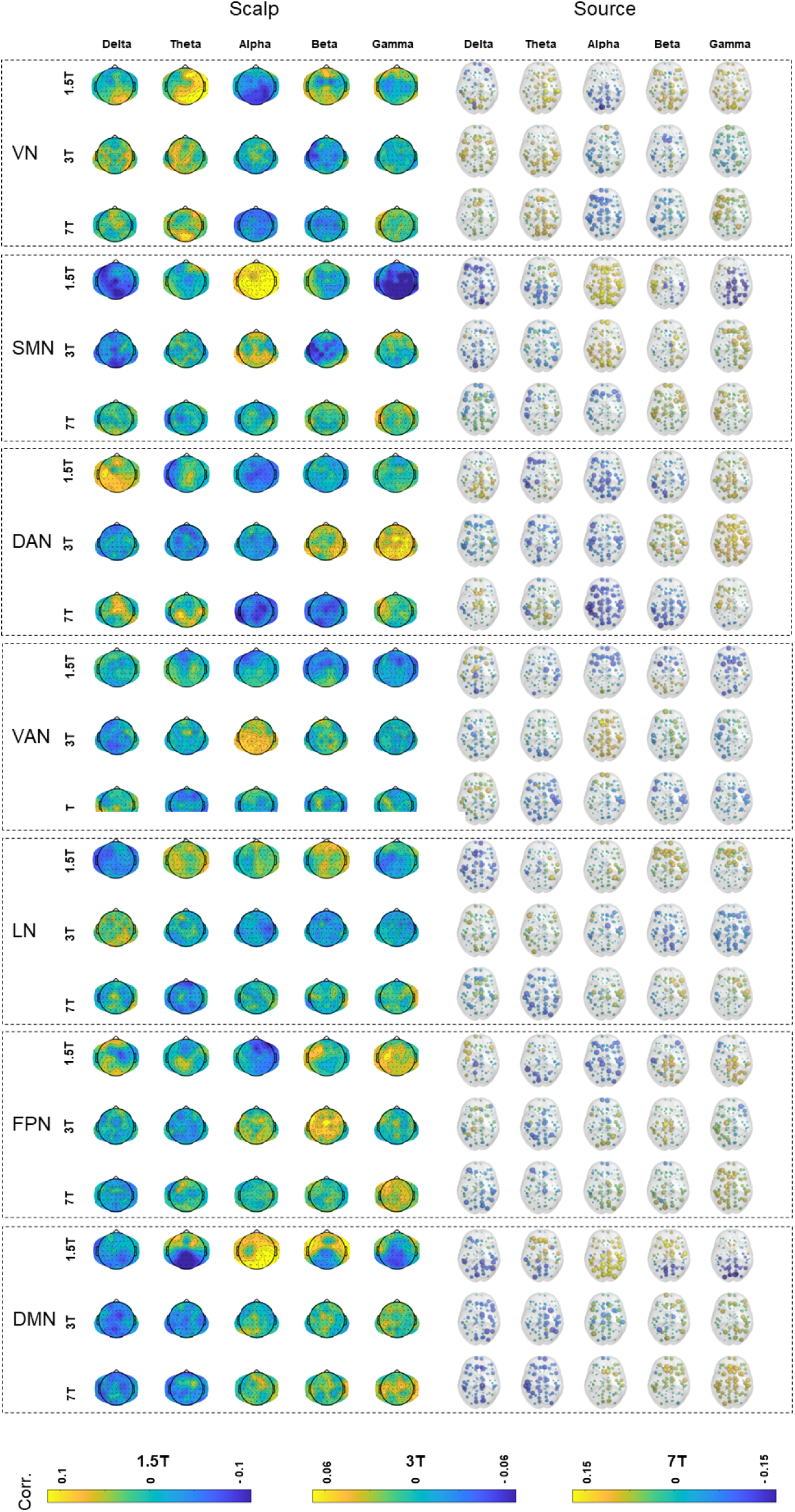
Temporal correlation spatial maps of EEG band-power and fMRI RSNs across different acquisition datasets for 10s HRF delay. The figure illustrates the subject-averaged spatial maps of the temporal correlations between 7 canonical fMRI resting state networks (RSNs) and EEG band-power across delta, theta, alpha, beta, and gamma bands for three independently acquired datasets (1.5T, 3T, and 7T), with EEG time-series convolved with an HRF with a 10-s overshoot delay. The left panel displays spatial maps related to EEG scalp data, while the right panel corresponds to source data, mapped to the Desikan-Killiany atlas.

### Spatially Averaged EEG-fMRI Correlations

**Fig. S7.**
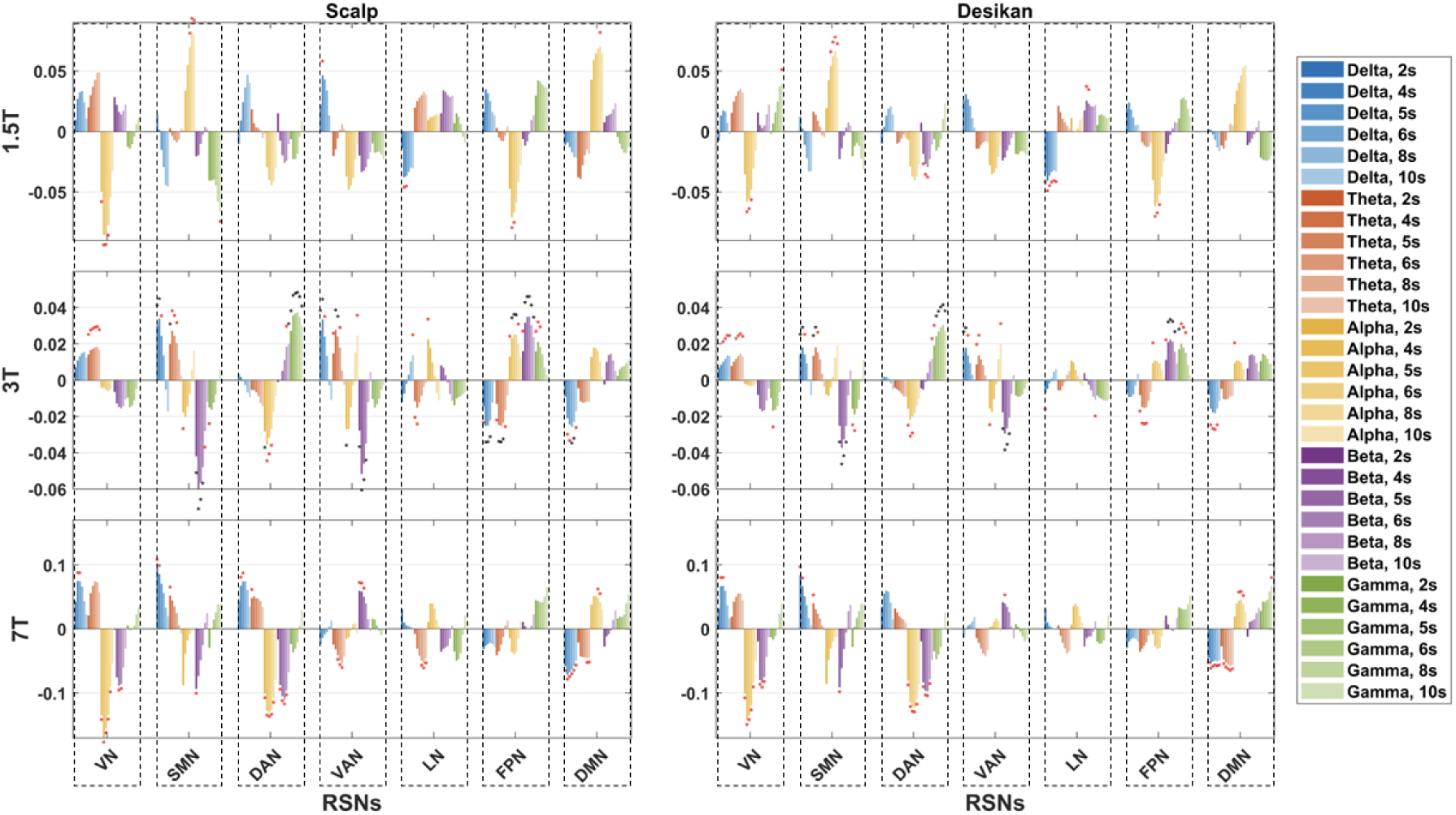
Spatially Averaged EEG-fMRI Correlations. The bar plots depict the spatially averaged temporal correlations between EEG band-power across five frequency-bands (delta, theta, alpha, beta, and gamma) and seven canonical fMRI Resting State Networks (RSNs), averaged across subjects for three independently acquired datasets (1.5T, 3T, and 7T). EEG time-series were convolved with a Hemodynamic Response Function (HRF) incorporating 2-s, 4-s, 5-s, 6-s, 8-s, and 10-s overshoot delays. On the left: EEG scalp data; on the right: EEG source (Desikan-Killiany atlas) data. Asterisks denote correlations significantly different from zero at p<0.05, as determined by a t-test across subjects; red and black asterisks represent uncorrected and False Discovery Rate-corrected results, respectively.

### Effect of the Number of Subjects on EEG-fMRI Correlations

**Fig. S8.**
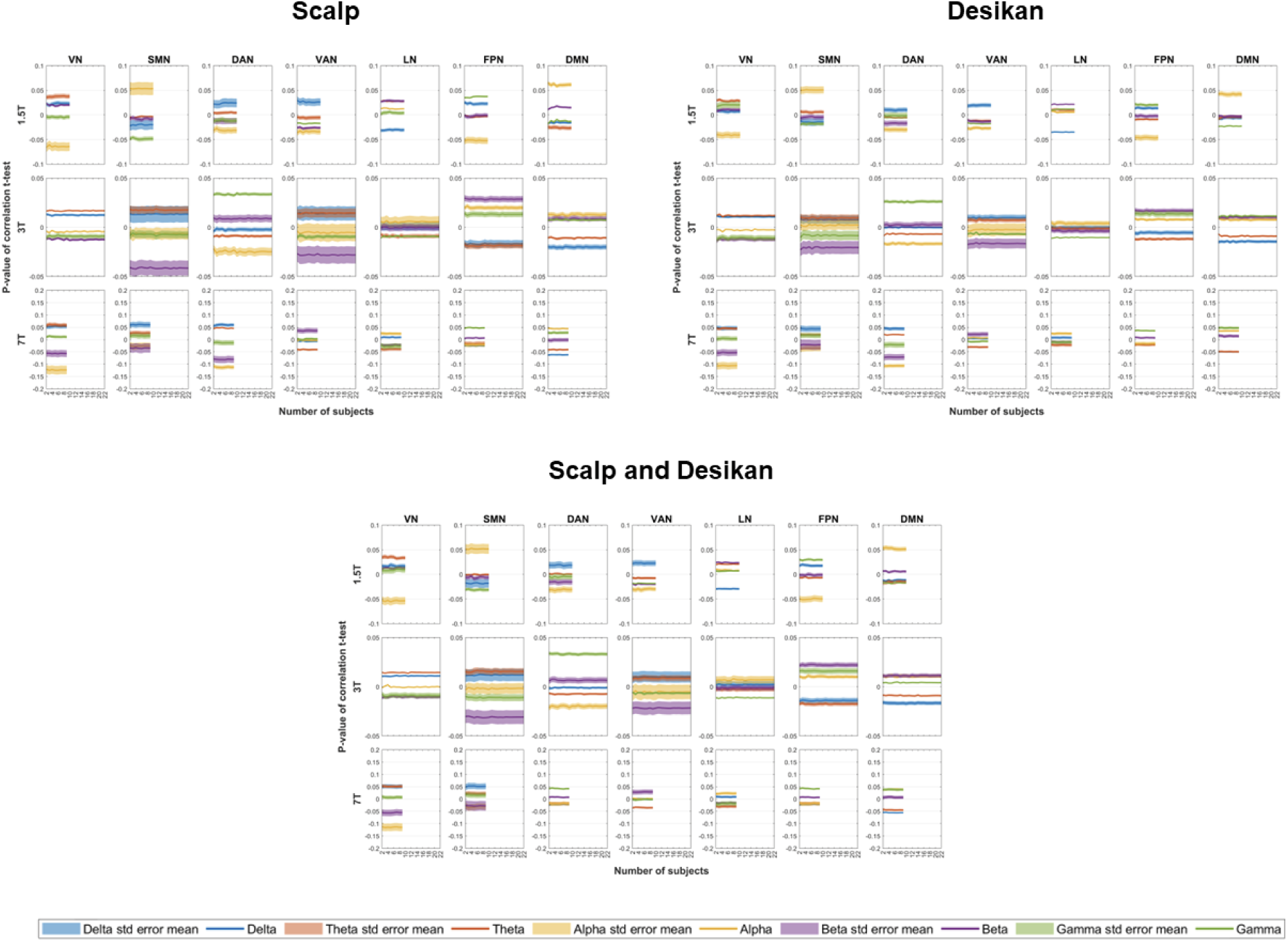
Effect of the number of subjects on EEG-fMRI correlations. Impact of increasing number of subjects on the correlation (averaged across subjects) derived from EEG-fMRI spatially averaged temporal correlations. Top left panel: EEG scalp data; Top right panel: EEG source (Desikan-Killiany atlas) data; Bottom panel: data pooled from both scalp and source (Desikan-Killiany atlas) EEG spaces to derive average correlation values. Distinct colors denote EEG band-power across delta, theta, alpha, beta, and gamma bands, with shaded areas indicating the standard mean error across a set of HRF delays: 2s, 4s, 5s, 6s, 8s, and 10s. Rows represent t-statistics calculated across subjects for each EEG-fMRI dataset (labeled as 1.5T, 3T, and 7T), while columns correspond to the seven canonical fMRI RSNs. For each dataset, subjects were randomly sampled (ranging from 1 to n subjects, over 5000 iterations) prior to computing the average temporal correlations values.

**Fig. S9.**
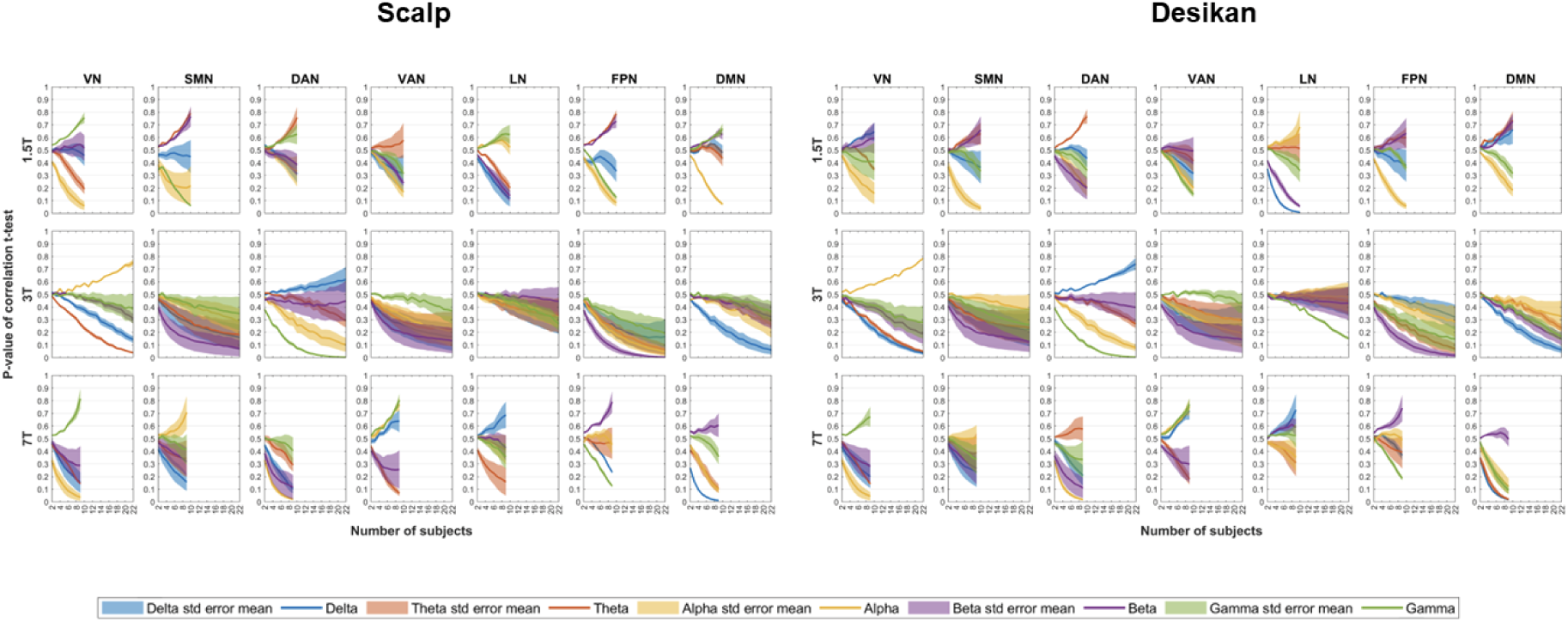
Effect of the number of subjects on the significance of EEG-fMRI correlations. Impact of increasing number of subjects on the p-value of the t-statistics derived from EEG-fMRI spatially averaged temporal correlations. On the left: EEG scalp data; on the right: EEG source (Desikan-Killiany atlas) data. Distinct colors denote EEG band-power across delta, theta, alpha, beta, and gamma bands, with shaded areas indicating the standard mean error across a set of HRF delays: 2s, 4s, 5s, 6s, 8s, and 10s. Rows represent t-statistics calculated across subjects for each EEG-fMRI dataset (labeled as 1.5T, 3T, and 7T), while columns correspond to the seven canonical fMRI RSNs. For each dataset, subjects were randomly sampled (ranging from 1 to n subjects, over 5000 iterations) prior to computing the t-stat values.

### Effect of the Scan Duration on EEG-fMRI Correlations

**Fig. S10.**
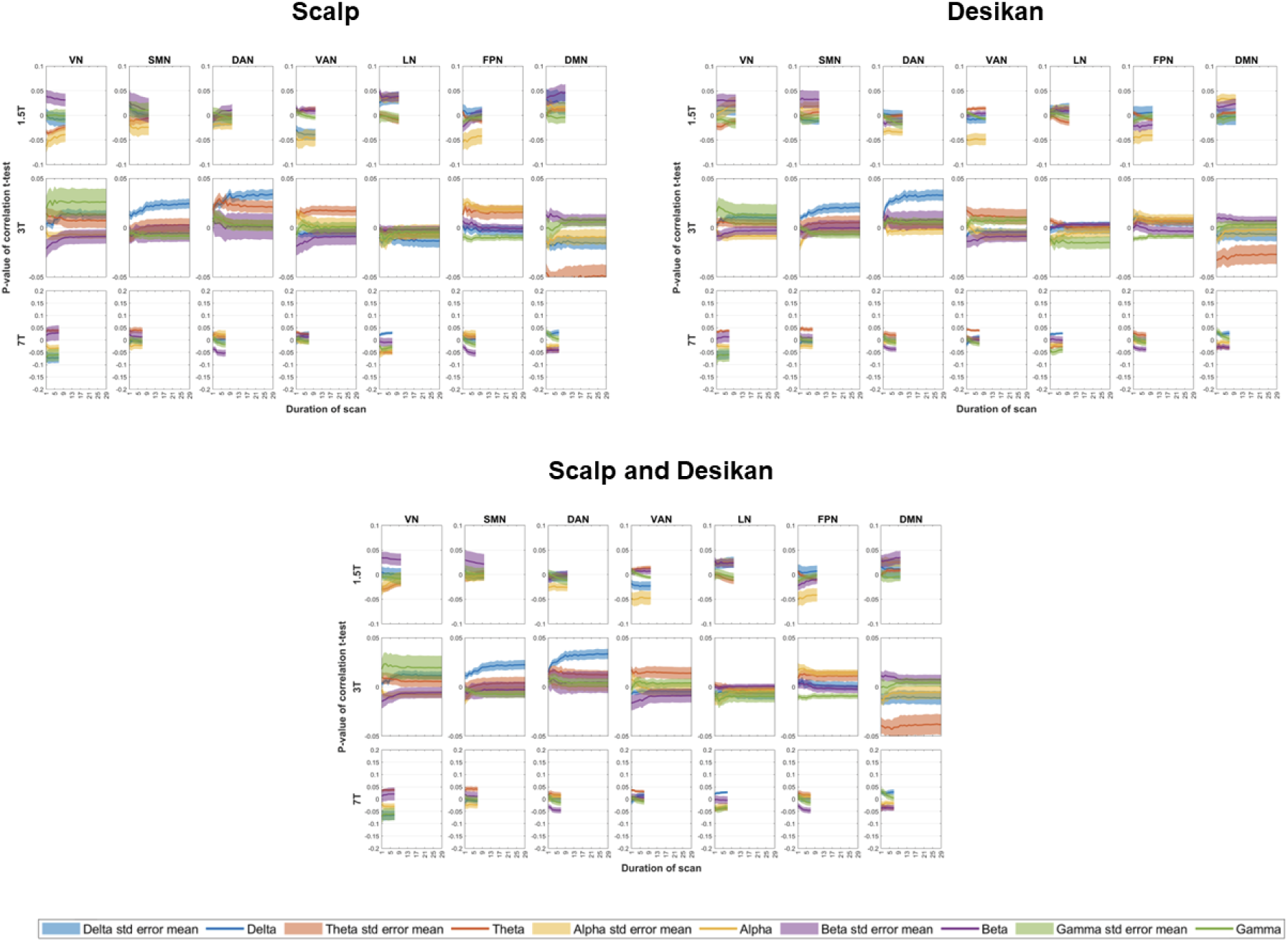
Effect of the scan duration on EEG-fMRI correlations. Impact of increasing scan duration on the correlation (averaged across subjects) derived from EEG-fMRI spatially averaged temporal correlations. Top left panel: EEG scalp data; Top right panel: EEG source (Desikan-Killiany atlas) data; Bottom panel: data pooled from both scalp and source (Desikan-Killiany atlas) EEG spaces to derive average correlation values. Distinct colors denote EEG band-power across delta, theta, alpha, beta, and gamma bands, with shaded areas indicating the standard mean error across a set of HRF delays: 2s, 4s, 5s, 6s, 8s, and 10s. Rows represent t-statistics calculated across subjects for each EEG-fMRI dataset (labeled as 1.5T, 3T, and 7T), while columns correspond to the seven canonical fMRI RSNs. For each dataset, segments of data were randomly selected (ranging from 1 to n consecutive minutes, over 5000 iterations) prior to computing the average temporal correlations values.

**Fig. S11.**
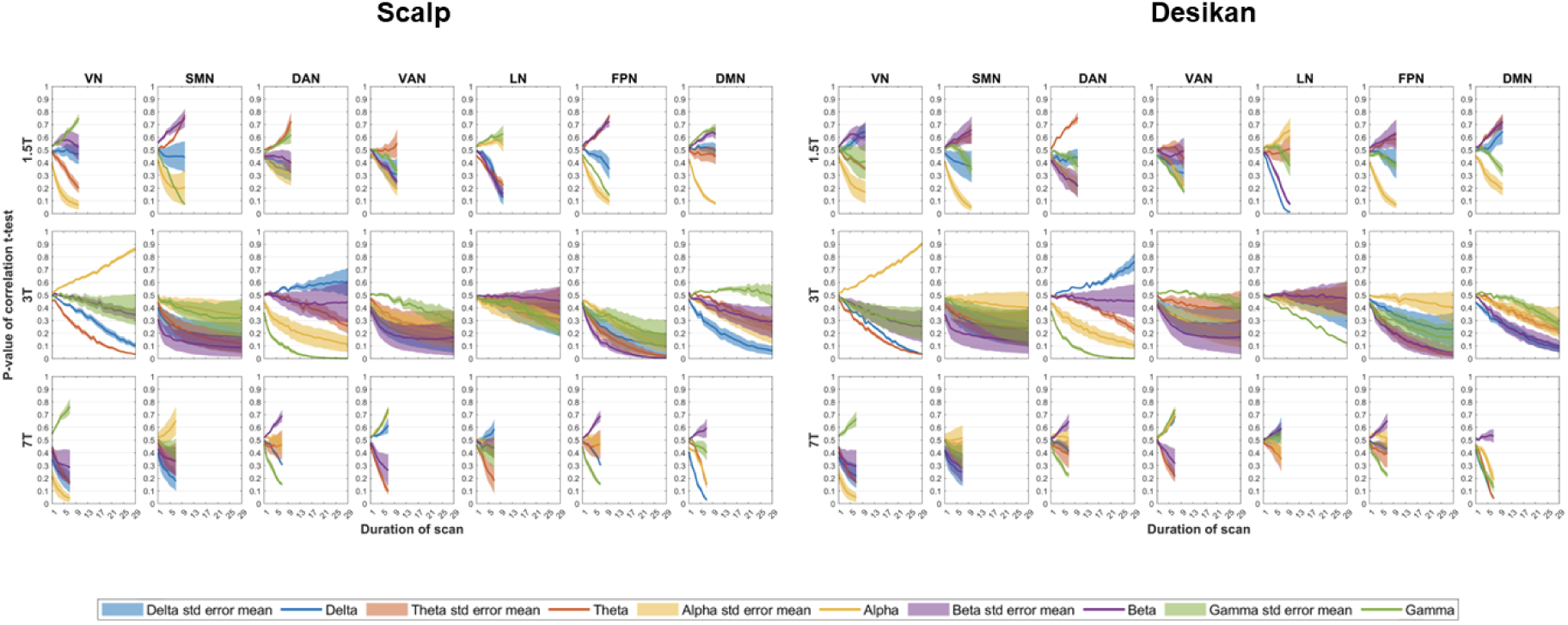
Effect of the scan duration on the significance of EEG-fMRI correlations. Impact of increasing scan duration on the p-value of the t-statistics derived from EEG-fMRI spatially averaged temporal correlations. On the left: EEG scalp data; on the right: EEG source (Desikan-Killiany atlas) data. Distinct colors denote EEG band-power across delta, theta, alpha, beta, and gamma bands, with shaded areas indicating the standard mean error across a set of HRF delays: 2s, 4s, 5s, 6s, 8s, and 10s. Rows represent t-statistics calculated across subjects for each EEG-fMRI dataset (labeled as 1.5T, 3T, and 7T), while columns correspond to the seven canonical fMRI RSNs. For each dataset, segments of data were randomly selected (ranging from 1 to n consecutive minutes, over 5000 iterations) prior to computing the temporal correlations and t-stat values.

### EEG-fMRI Correlations Across Extended HRF delays

**Fig. S12.**
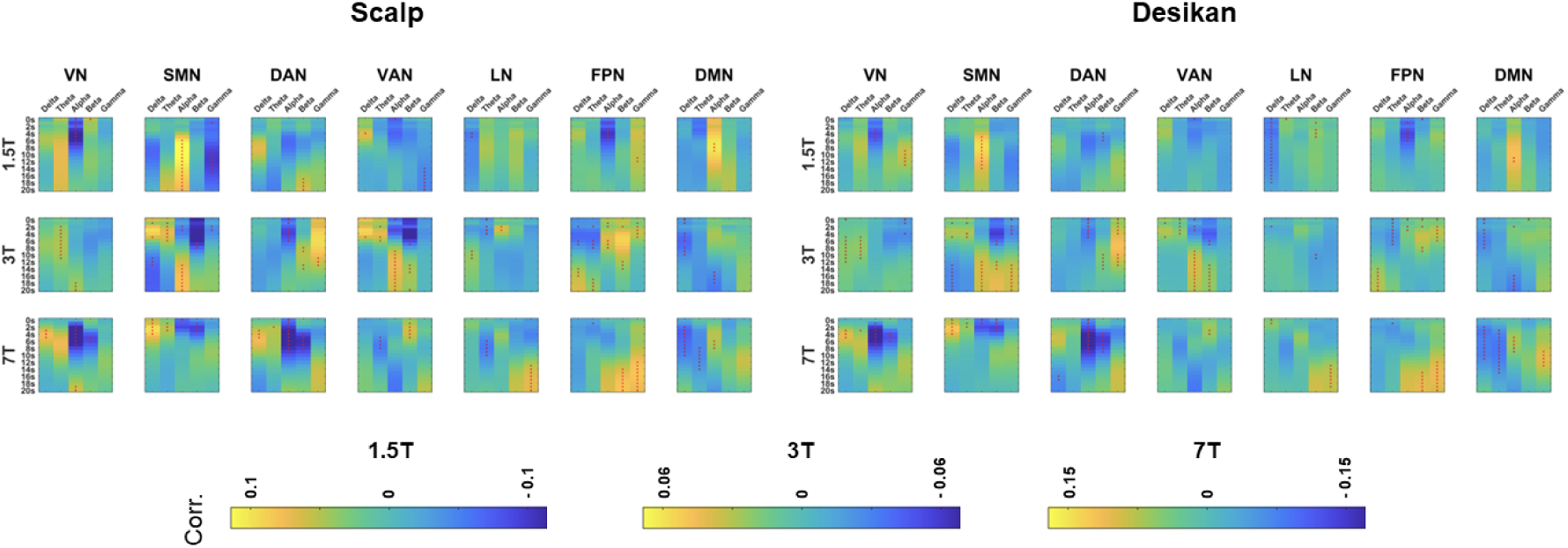
EEG-fMRI correlations across an extended range of HRF delays. The heatmaps display the value of the spatially averaged EEG-fMRI correlations, further averaged across subjects, for different Hemodynamic Response Function (HRF) delays, ranging from 0 to 20s. Each subplot corresponds to a combination of the independently acquired datasets (1.5T, 3T and 7T) and 7 canonical Resting State Network (RSN). On the left: EEG scalp data; on the right: EEG source (Desikan-Killiany atlas) data. Red dots denote correlation values that are significantly different from zero, as per t-tests (uncorrected for multiple comparisons). In each subplot, the correlation values corresponding to increasing HRF delays are displayed from top to bottom, and the values corresponding to each EEG frequency-band are displayed from left to right.

## References

Abreu, R., Jorge, J., Leal, A., Koenig, T., & Figueiredo, P. (2021). EEG Microstates Predict Concurrent fMRI Dynamic Functional Connectivity States. Brain Topography, 34(1), 41–55. 10.1007/s10548-020-00805-1

Abreu, R., Leal, A., & Figueiredo, P. (2018). EEG-informed fMRI: A review of data analysis methods. Frontiers in Human Neuroscience, 12(February), 1–23. 10.3389/fnhum.2018.00029

Allen, E. A. (2018). EEG Signatures of Dynamic Functional Network Connectivity. Brain Topography, 31(1), 101–116. 10.1007/s10548-017-0546-2

Andersson, J. L. R., Jenkinson, M., & Smith, S. (2007). Non-linear registration aka Spatial normalisation FMRIB Technial Report TR07JA2. FMRIB Technial Report TR07JA2.

Andersson, J. L. R., Skare, S., & Ashburner, J. (2003). How to correct susceptibility distortions in spin-echo echo-planar images: application to diffusion tensor imaging. NeuroImage, 20, 870–888. 10.1016/S1053-8119(03)00336-7

Baillet, S., Mosher, J. C., Pantazis, D., & Leahy, R. M. (2011). Brainstorm : A User-Friendly Application for MEG / EEG Analysis Franc. Computational Intelligence and Neuroscience, 2011. 10.1155/2011/879716

Beckmann, C. F., & Smith, S. M. (2004). Probabilistic independent component analysis for functional magnetic resonance imaging. IEEE transactions on medical imaging, 23(2), 137–152. 10.1109/TMI.2003.822821

Biswal, B., Yetkin, F. Z., Haughton, V. M., & Hyde, J. S. (1995). Functional connectivity in the motor cortex of resting human brain using echo-planar MRI. Magnetic resonance in medicine, 34(4), 537–541. 10.1002/mrm.1910340409

Bowman, A. D., Grif, J. C., Visscher, K. M., Dobbins, A. C., Gawne, T. J., Difrancesco, M. W., & Sza, J. P. (2017). Relationship Between Alpha Rhythm and the Default Mode Network : Journal of Clinical Neurophysiology, 0(0). 10.1097/WNP.0000000000000411

Braga, R. M., & Buckner, R. L. (2017). Parallel Interdigitated Distributed Networks within the Individual Estimated by Intrinsic Functional Connectivity. Neuron, 95(2), 457–471.e5. 10.1016/j.neuron.2017.06.038

Britz, J., Ville, D. Van De, & Michel, C. M. (2010). NeuroImage BOLD correlates of EEG topography reveal rapid resting-state network dynamics. NeuroImage, 52(4), 1162–1170. 10.1016/j.neuroimage.2010.02.052

Caplan, J. B., Madsen, J. R., Schulze-Bonhage, A., Aschenbrenner-Scheibe, R., Newman, E. L., & Kahana, M. J. (2003). Human Oscillations Related to Sensorimotor Integration and Spatial Learning.

Cavanagh, J. F., & Frank, M. J. (2014). Frontal theta as a mechanism for cognitive control. In Trends in Cognitive Sciences (Vol. 18, Issue 8, pp. 414–421). Elsevier Ltd. 10.1016/j.tics.2014.04.012

Chang, C., Cunningham, J. P., & Glover, G. H. (2009). NeuroImage Influence of heart rate on the BOLD signal : The cardiac response function. NeuroImage, 44(3), 857–869. 10.1016/j.neuroimage.2008.09.029

Chang, C., Liu, Z., Chen, M. C., Liu, X., & Duyn, J. H. (2013). NeuroImage EEG correlates of time-varying BOLD functional connectivity. NeuroImage, 72, 227–236. 10.1016/j.neuroimage.2013.01.049

Chang, C., & Glover, G. H. (2010). Time-frequency dynamics of resting-state brain connectivity measured with fMRI. NeuroImage, 50(1), 81–98. 10.1016/j.neuroimage.2009.12.011

Chang, C., & Chen, J. E. (2021). Multimodal EEG-fMRI: advancing insight into large-scale human brain dynamics. Current opinion in biomedical engineering, 18, 100279. 10.1016/j.cobme.2021.100279

Collins, D. L., Neelin, P., Peters, T. M., & Evans, A. C. (1994). Automatic 3D intersubject registration of MR volumetric data in standardized Talairach space. Journal of computer assisted tomography, 18(2), 192–205.

Desikan, R. S., Ségonne, F., Fischl, B., Quinn, B. T., Dickerson, B. C., Blacker, D., Buckner, R. L., Dale, A. M., Maguire, R. P., Hyman, B. T., Albert, M. S., & Killiany, R. J. (2006). An automated labeling system for subdividing the human cerebral cortex on MRI scans into gyral based regions of interest. NeuroImage, 31(3), 968–980.

Deligianni, F., Centeno, M., Carmichael, D. W., & Clayden, J. D. (2014). Relating resting-state fMRI and EEG whole-brain connectomes across frequency-bands. Frontiers in Neuroscience, 8(August). 10.3389/fnins.2014.00258

Dice, L. R. (1945). Measures of the Amount of Ecologic Association Between Species. Ecological Society of America, 26(3), 297–302.

DiFrancesco, M. W. (2009). Simultaneous EEG/Functional Magnetic Resonance Imaging at 4 Tesla: Correlates of Brain Activity to Spontaneous Alpha Rhythm During Relaxation. Journal of Clinical Neurophysiology, 25(April 2005), 255–264. 10.1097/WNP.0b013e3181879d56.Simultaneous

Dixon, M. L., de La Vega, A., Mills, C., Andrews-Hanna, J., Spreng, R. N., Cole, M. W., & Christoff, K. (2018). Heterogeneity within the frontoparietal control network and its relationship to the default and dorsal attention networks. Proceedings of the National Academy of Sciences of the United States of America, 115(7), E1598–E1607. 10.1073/pnas.1715766115

Dosenbach, N. U., Fair, D. A., Miezin, F. M., Cohen, A. L., Wenger, K. K., Dosenbach, R. A., Fox, M. D., Snyder, A. Z., Vincent, J. L., Raichle, M. E., Schlaggar, B. L., & Petersen, S. E. (2007). Distinct brain networks for adaptive and stable task control in humans. Proceedings of the National Academy of Sciences of the United States of America, 104(26), 11073–11078. 10.1073/pnas.0704320104

Falahpour, M., Chang, C., Wong, C. W., & Liu, T. T. (2018). Template-based prediction of vigilance fluctuations in resting-state fMRI. NeuroImage, 174, 317–327. 10.1016/j.neuroimage.2018.03.012

Fleury, M., Figueiredo, P., Vourvopoulos, A., & Lécuyer, A. (2023). Two is better? combining EEG and fMRI for BCI and neurofeedback: a systematic review. Journal of neural engineering, 20(5), 10.1088/1741-2552/ad06e1. 10.1088/1741-2552/ad06e1

Fischl, B. (2012). FreeSurfer. NeuroImage FreeSurfer. NeuroImage, 62, 774–781. 10.1016/j.neuroimage.2012.01.021

Fox, M. D., Corbetta, M., Snyder, A. Z., Vincent, J. L., & Raichle, M. E. (2006). Spontaneous neuronal activity distinguishes human dorsal and ventral attention systems. www.pnas.orgcgidoi10.1073pnas.0604187103

Glover GH, Li TQ, Ress D. (2000) Image-based method for retrospective correction of physiological motion effects in fMRI: RETROICOR. Magn Reson Med. Jul;44(1):162–7. https://doi.org.10.1002/1522-2594(200007)44:1<162::aid-mrm23>3.0.co;2-e

Goldman, R. I. (2002). Simultaneous EEG and fMRI of the alpha rhythm. NeuroReport, 13(18), 2487–2492. 10.1097/01.wnr.0000047685.08940.d0

Gonçalves, S. I., de Munck, J. C., Pouwels, P. J., Schoonhoven, R., Kuijer, J. P., Maurits, N. M., Hoogduin, J. M., Van Someren, E. J., Heethaar, R. M., & Lopes da Silva, F. H. (2006). Correlating the alpha rhythm to BOLD using simultaneous EEG/fMRI: inter-subject variability. NeuroImage, 30(1), 203–213. 10.1016/j.neuroimage.2005.09.062

Gramfort, A., Papadopoulo, T., Olivi, E., Clerc, M., 2010. OpenMEEG: opensource software for quasistatic bioelectromagnetics. Biomed. Eng. OnLine 9, 45. 10.1186/1475-925X-9-45

Greicius, M. D., Krasnow, B., Reiss, A. L., Menon, V., & Raichle, M. E. (n.d.). Functional connectivity in the resting brain: A network analysis of the default mode hypothesis. www.pnas.org.

Hutchison, R. M., Womelsdorf, T., Allen, E. A., Bandettini, P. A., Calhoun, V. D., Corbetta, M., Della, S., Duyn, J. H., Glover, G. H., Gonzalez-castillo, J., Handwerker, D. A., Keilholz, S., Kiviniemi, V., Leopold, D. A., Pasquale, F. De, Sporns, O., Walter, M., & Chang, C. (2013). NeuroImage Dynamic functional connectivity : Promise, issues, and interpretations. NeuroImage, 80, 360–378. 10.1016/j.neuroimage.2013.05.079

Jann, K., Kottlow, M., Dierks, T., Boesch, C., & Koenig, T. (2010). Topographic Electrophysiological Signatures of fMRI Resting State Networks Definition of frequency-bands. PLoS ONE, 5(9). 10.1371/journal.pone.0012945

Jenkinson, M., & Smith, S. (2001). A global optimisation method for robust affine registration of brain images. Medical Image Analysis, 5, 143–156.

Jenkinson, M., Bannister, P., Brady, M., & Smith, S. (2002). Improved Optimization for the Robust and Accurate Linear Registration and Motion Correction of Brain Images. NeuroImage, 841, 825–841. 10.1006/nimg.2002.1132

Joliot, M., Cremona, S., Tzourio, C., & Etard, O. (2024). Modulate the impact of the drowsiness on the resting state functional connectivity. Scientific reports, 14(1), 8652. 10.1038/s41598-024-59476-8

Jorge, J., Bouloc, C., Bréchet, L., Michel, C. M., & Gruetter, R. (2019). Investigating the variability of cardiac pulse artifacts across heartbeats in simultaneous EEG-fMRI recordings: A 7T study. NeuroImage, 191(February), 21–35. 10.1016/j.neuroimage.2019.02.021

Jorge, J., Grouiller, F., Ipek, Ö., Stoermer, R., Michel, C. M., Figueiredo, P., Zwaag, W. Van Der, & Gruetter, R. (2015). NeuroImage Simultaneous EEG – fMRI at ultra-high field : Artifact prevention and safety assessment. NeuroImage, 105, 132–144. 10.1016/j.neuroimage.2014.10.055

Jorge, J., van der Zwaag, W., & Figueiredo, P. (2014). EEG-fMRI integration for the study of human brain function. NeuroImage, 102 Pt 1, 24–34. 10.1016/j.neuroimage.2013.05.114

Keilholz, S., Caballero-Gaudes, C., Bandettini, P., Deco, G., & Calhoun, V. (2017). Time-Resolved Resting-State Functional Magnetic Resonance Imaging Analysis: Current Status, Challenges, and New Directions. Brain connectivity, 7(8), 465–481. 10.1089/brain.2017.0543

Kybic, J., Clerc, M., Abboud, T., Faugeras, O., Keriven, R., Papadopoulo, T., 2005. A common formalism for the Integral formulations of the forward EEG problem. IEEE Trans. Med. Imaging 24, 12–28. 10.1109/TMI.2004.837363

Labounek, R., Bridwell, D. A., Mareček, R., Lamoš, M., Mikl, M., Bednařík, P., Baštinec, J., Slavíček, T., Hluštík, P., Brázdil, M., & Jan, J. (2019). EEG spatiospectral patterns and their link to fMRI BOLD signal via variable hemodynamic response functions. Journal of Neuroscience Methods, 318(February), 34–46. 10.1016/j.jneumeth.2019.02.012

Laufs, H., Holt, J. L., Elfont, R., Krams, M., Paul, J. S., Krakow, K., & Kleinschmidt, A. (2006). Where the BOLD signal goes when alpha EEG leaves. NeuroImage. Clinical, 31, 1408–1418. 10.1016/j.neuroimage.2006.02.002

Laufs, H., Kleinschmidt, A., Beyerle, A., Eger, E., Salek-haddadi, A., Preibisch, C., & Krakow, K. (2003). EEG-correlated fMRI of human alpha activity. NeuroImage, 19, 1463–1476. 10.1016/S1053-8119(03)00286-6

Laufs, H., Krakow, K., Sterzer, P., Eger, E., Beyerle, A., & Kleinschmidt, A. (2003). Electroencephalographic signatures of attentional and cognitive default modes in spontaneous brain activity fluctuations at rest. PNAS, 100(19).

Logothetis, N., Pauls, J., Augath, M. et al. (2001) Neurophysiological investigation of the basis of the fMRI signal. Nature 412, 150–157. 10.1038/35084005

Logothetis, N. K., & Wandell, B. A. (2004). Interpreting the BOLD signal. Annual review of physiology, 66, 735–769. 10.1146/annurev.physiol.66.082602.092845

Magosso, E., Ricci, G., & Ursino, M. (2021). Alpha and theta mechanisms operating in internal-external attention competition. Journal of Integrative Neuroscience, 20(1), 1–19. 10.31083/J.JIN.2021.01.422

Makeig, S., & Jung, T. P. (1995). Changes in alertness are a principal component of variance in the EEG spectrum. Neuroreport, 7(1), 213–216.

Mantini, D., Perrucci, M. G., Del Gratta, C., Romani, G. L., & Corbetta, M. (2007). Electrophysiological signatures of resting state networks in the human brain. Proceedings of the National Academy of Sciences of the United States of America, 104(32), 13170–13175. 10.1073/pnas.0700668104

Marawar, R. A., Yeh, H. J., Carnabatu, C. J., & Stern, J. M. (2017). Functional MRI Correlates of Resting-State Temporal Theta and Delta EEG Rhythms. Journal of Clinical Neurophysiology, 34(1). 10.1097/WNP.0000000000000309

Mayhew, S. D., Ostwald, D., Porcaro, C., & Bagshaw, A. P. (2013). Spontaneous EEG alpha oscillation interacts with positive and negative BOLD responses in the visual-auditory cortices and default-mode network. NeuroImage, 76, 362–372. 10.1016/j.neuroimage.2013.02.070

Meir-Hasson, Y., Kinreich, S., Podlipsky, I., Hendler, T., & Intrator, N. (2014). NeuroImage An EEG Finger-Print of fMRI deep regional activation. NeuroImage, 102, 128–141. 10.1016/j.neuroimage.2013.11.004

Meyer, M. C., Oort, E. S. B. Van, & Barth, M. (2013). Electrophysiological Correlation Patterns of Resting State Networks in Single Subjects : A Combined EEG – fMRI Study. Brain Topography, 98–109. 10.1007/s10548-012-0235-0

Mo, J., Liu, Y., Huang, H., & Ding, M. (2013). NeuroImage Coupling between visual alpha oscillations and default mode activity ☆. NeuroImage, 68, 112–118. 10.1016/j.neuroimage.2012.11.058

Moosmann, M., Ritter, P., Krastel, I., Brink, A., Thees, S., Blankenburg, F., Taskin, B., Obrig, H., & Villringer, A. (2003). Correlates of alpha rhythm in functional magnetic resonance imaging and near infrared spectroscopy. NeuroImage, 20, 145–158. 10.1016/S1053-8119(03)00344-6

Morillon, B., Lehongre, K., Frackowiak, R. S., Ducorps, A., Kleinschmidt, A., Poeppel, D., & Giraud, A. L. (2010). Neurophysiological origin of human brain asymmetry for speech and language. Proceedings of the National Academy of Sciences of the United States of America, 107(43), 18688–18693. 10.1073/pnas.1007189107

Musso, F., Brinkmeyer, J., Mobascher, A., Warbrick, T., & Winterer, G. (2010). NeuroImage Spontaneous brain activity and EEG microstates. A novel EEG / fMRI analysis approach to explore resting-state networks. NeuroImage, 52(4), 1149–1161. 10.1016/j.neuroimage.2010.01.093

Nickerson, L. D., Smith, S. M., Öngür, D., & Beckmann, C. F. (2017). Using Dual Regression to Investigate Network Shape and Amplitude in Functional Connectivity Analyses. Frontiers in Neuroscience, 11(March), 1–18. 10.3389/fnins.2017.00115

Penny, William & Friston, Karl & Ashburner, John & Kiebel, Stefan & Nichols, T.. (2007). Statistical Parametric Mapping: The Analysis of Functional Brain Images. 10.1016/B978-0-12-372560-8.X5000-1.

Portnova, G. V, Tetereva, A., Balaev, V., & Atanov, M. (2018). Correlation of BOLD Signal with Linear and Nonlinear Patterns of EEG in Resting State EEG-Informed fMRI. Frontiers in Human Neuroscience, 11(January). 10.3389/fnhum.2017.00654

Raichle, M. E., Macleod, A. M., Snyder, A. Z., Powers, W. J., Gusnard, D. A., & Shulman, G. L. (2001). A default mode of brain function. Proc Natl Acad Sci U S A., 98(2), 676–682. 10.1073/pnas.98.2.676

Rajkumar, R., Lerche, C., Shah, N. J., Farrher, E., Mauler, J., Sripad, P., Brambilla, C. R., Kops, E. R., Langen, K., Neuner, I., Scheins, J., & Dammers, J. (2021). Comparison of EEG microstates with resting state fMRI and FDG-PET measures in the default mode network via simultaneously recorded trimodal ( PET / MR / EEG ) data. May 2018, 4122–4133. 10.1002/hbm.24429

Ritter, P., Moosmann, M., & Villringer, A. (2009). Rolandic alpha and beta EEG rhythms’ strengths are inversely related to fMRI-BOLD signal in primary somatosensory and motor cortex. Human Brain Mapping, 30(4), 1168–1187. 10.1002/hbm.20585

Sadaghiani, S., Lehongre, K., Morillon, B., & Giraud, A. (2010). Intrinsic Connectivity Networks, Alpha Oscillations, and Tonic Alertness : A Simultaneous Electroencephalography / Functional Magnetic Resonance Imaging Study. The Journal of Neuroscience, 30(30), 10243–10250. 10.1523/JNEUROSCI.1004-10.2010

Scheeringa, R., Bastiaansen, M. C. M., Magnus, K., Oostenveld, R., Norris, D. G., & Hagoort, P. (2008). Frontal theta EEG activity correlates negatively with the default mode network in resting state. International Journal of Psychophysiology, 67, 242–251. 10.1016/j.ijpsycho.2007.05.017

Seeley, W. W., Menon, V., Schatzberg, A. F., Keller, J., Glover, G. H., Kenna, H., Reiss, A. L., & Greicius, M. D. (2007). Dissociable intrinsic connectivity networks for salience processing and executive control. The Journal of neuroscience : the official journal of the Society for Neuroscience, 27(9), 2349–2356. 10.1523/JNEUROSCI.5587-06.2007

Simões, M., Abreu, R., Direito, B., Sayal, A., Castelhano, J., Carvalho, P., & Castelo-Branco, M. (2020). How much of the BOLD-fMRI signal can be approximated from simultaneous EEG data: relevance for the transfer and dissemination of neurofeedback interventions. Journal of neural engineering, 17(4), 046007. 10.1088/1741-2552/ab9a98

Smith, S. M. (2002). Fast Robust Automated Brain Extraction. Human Brain Mapping, 155, 143–155. 10.1002/hbm.10062

Smith, S. M., & Brady, J. M. (1997). SUSAN — A New Approach to Low Level Image Processing. International Journal of Computer Vision, 23(1), 45–78.

Smith, S. M., Fox, P. T., Miller, K. L., Glahn, D. C., Fox, P. M., Mackay, C. E., Filippini, N., Watkins, K. E., Toro, R., Laird, A. R., & Beckmann, C. F. (2009). Correspondence of the brain’s functional architecture during activation and rest. Proceedings of the National Academy of Sciences of the United States of America, 106(31), 13040–13045. 10.1073/pnas.0905267106

Smith, S. M., Jenkinson, M., Woolrich, M. W., Beckmann, C. F., Behrens, T. E. J., Johansen-berg, H., Bannister, P. R., Luca, M. De, Drobnjak, I., Flitney, D. E., Niazy, R. K., Saunders, J., Vickers, J., Zhang, Y., Stefano, N. De, Brady, J. M., & Matthews, P. M. (2004). Advances in functional and structural MR image analysis and implementation as FSL. NeuroImage, 23, 208–219. 10.1016/j.neuroimage.2004.07.051

Spreng, R. N., Sepulcre, J., Turner, G. R., Stevens, W. D., & Schacter, D. L. (2013). Intrinsic architecture underlying the relations among the default, dorsal attention, and frontoparietal control networks of the human brain. Journal of Cognitive Neuroscience, 25(1), 74–86. 10.1162/jocn_a_00281

Stephani, C. (2014). Limbic System. In Encyclopedia of the Neurological Sciences (pp. 897–900). Elsevier Inc. 10.1016/B978-0-12-385157-4.01157-X

Tagliazucchi, E., Wegner, F. Von, Morzelewski, A., Brodbeck, V., & Laufs, H. (2012). Dynamic BOLD functional connectivity in humans and its electrophysiological correlates. 6(December), 1–23. 10.3389/fnhum.2012.00339

Tagliazucchi, E., & Laufs, H. (2014). Decoding wakefulness levels from typical fMRI resting-state data reveals reliable drifts between wakefulness and sleep. Neuron, 82(3), 695–708. 10.1016/j.neuron.2014.03.020

Tsuchimoto, S., Shibusawa, S., Mizuguchi, N., Kato, K., Ebata, H., Liu, M., Hanakawa, T., & Ushiba, J. (2017). Resting-State Fluctuations of EEG Sensorimotor Rhythm Reflect BOLD Activities in the Pericentral Areas: A Simultaneous EEG-fMRI Study. Frontiers in Human Neuroscience, 11. 10.3389/FNHUM.2017.00356

Uddin, L. Q., Betzel, R. F., Cohen, J. R., Damoiseaux, J. S., De Brigard, F., Eickhoff, S. B., Fornito, A., Gratton, C., Gordon, E. M., Laird, A. R., Larson-Prior, L., McIntosh, A. R., Nickerson, L. D., Pessoa, L., Pinho, A. L., Poldrack, R. A., Razi, A., Sadaghiani, S., Shine, J. M., … Spreng, R. N. (2023). Controversies and progress on standardization of large-scale brain network nomenclature. Network Neuroscience, 7(3), 864–905. 10.1162/netn_a_00323

Vincent, J. L., Kahn, I., Snyder, A. Z., Raichle, M. E., & Buckner, R. L. (2008). Evidence for a frontoparietal control system revealed by intrinsic functional connectivity. Journal of Neurophysiology, 100(6), 3328–3342. 10.1152/jn.90355.2008

Wirsich, J., Giraud, A. L., & Sadaghiani, S. (2020). Concurrent EEG- and fMRI-derived functional connectomes exhibit linked dynamics. NeuroImage, 219(February), 116998. 10.1016/j.neuroimage.2020.116998

Wirsich, J., Jorge, J., Iannotti, G. R., Shamshiri, E. A., Grouiller, F., Abreu, R., Lazeyras, F., Giraud, A. L., Gruetter, R., Sadaghiani, S., & Vulliémoz, S. (2021). The relationship between EEG and fMRI connectomes is reproducible across simultaneous EEG-fMRI studies from 1.5T to 7T. NeuroImage, 231(February), 117864. 10.1016/j.neuroimage.2021.117864

Wirsich, J., Bagshaw, A., Guye, M., Lemieux, L., & Bénar, C.-G. (2023). Experimental design and data analysis strategies. In Springer. 10.1007/978-3-031-07121-8_12

Yeo, B. T. T., Krienen, F. M., Sepulcre, J., Sabuncu, M. R., Hollinshead, M., Roffman, J. L., Smoller, J. W., Polimeni, J. R., Fischl, B., Liu, H., Buckner, R. L., Kaiser, R. H., Andrews-hanna, J. R., Spielberg, J. M., Warren, S. L., Sutton, B. P., Miller, G. A., Heller, W., Banich, M. T., … Buckner, R. L. (2014). The organization of the human cerebral cortex estimated by intrinsic functional connectivity The organization of the human cerebral cortex estimated by intrinsic functional connectivity. Journal of Neurophysiology. 10.1152/jn.00338.2011

Yin, S., Liu, Y., & Ding, M. (2016). Amplitude of Sensorimotor Mu Rhythm Is Correlated with BOLD from Multiple Brain Regions : A Simultaneous EEG-fMRI Study. Frontiers in Human Neuroscience, 10(July), 1–12. 10.3389/fnhum.2016.00364

Yuan, S., Cohen, D. B., Ravel, J., Abdo, Z., & Forney, L. J. (2012). Evaluation of Methods for the Extraction and Purification of DNA from the Human Microbiome. 7(3). 10.1371/journal.pone.0033865

Zhang, Y., Brady, M., & Smith, S. (2001). Segmentation of Brain MR Images Through a Hidden Markov Random Field Model and the Expectation-Maximization Algorithm. IEEE Transactions on Biomedical Engineering, 20(1), 45–57.

